# Age-related differences in prestimulus subsequent memory effects assessed with event-related potentials

**DOI:** 10.1101/188664

**Authors:** Joshua D. Koen, Erin D. Horne, Nedra Hauck, Michael D. Rugg

## Abstract

Prestimulus subsequent memory effects (preSMEs) – differences in neural activity elicited by a task cue at encoding that are predictive of later memory performance – are thought to reflect differential engagement of preparatory processes that benefit episodic memory encoding. We investigated age differences in preSMEs indexed by differences in ERP amplitude just prior to the onset of a study item. Young and older adults incidentally encoded words for a subsequent memory test. Each study word was preceded by a task cue that signaled the judgment to perform on the word. Words were presented for either a short (300 ms) or a long (1000 ms) duration with the aim of placing differential benefits on engaging preparatory processes initiated by the task cue. ERPs associated with subsequent successful and unsuccessful recollection, operationalized here by source memory accuracy, were estimated time-locked to the onset of the task cue. In a late time-window (1000-2000 ms following onset of the cue), young adults demonstrated frontally distributed preSMEs for both the short and the long study durations, albeit with opposite polarities in the two conditions. This finding suggests that preSMEs in young adults are sensitive to perceived task demands. Although older adults showed no evidence of preSMEs in the same late time window, significant preSMEs were observed in an earlier time window (500-1000 ms) that were invariant with study duration. These results are broadly consistent with the proposal that older adults differ from their younger counterparts in how they engage preparatory processes during memory encoding.

## Introduction

Aging is associated with a decline in the ability to recollect specific details about prior events (for recent reviews, see Koen & Yonelinas, 2014; Schoemaker, Gauthier, & Pruessner, 2014). This decline is partly, if not largely, due to age-related reductions in the efficacy of encoding (Craik, 1983; Craik & Byrd, 1982; Craik & Rose, 2012; Friedman & Johnson, 2014; Werkle-Bergner, Muller, Li, & Lindenberger, 2006). This evidence has motivated numerous studies that have used fMRI or, less frequently, event-related potentials (ERPs), to identify the neural bases of age-related differences in the efficacy of encoding operations supporting later recollection. These studies have almost invariably employed some version of the ‘subsequent memory procedure’, which allows identification of differences in neural activity (‘subsequent memory effects’ or SMEs) elicited by study items that are remembered or forgotten on a later memory test (Wagner et al., 1998; for reviews, see Kim, 2011; Paller & Wagner, 2002; Rugg, Johnson, & Uncapher, 2015). The findings from these studies strongly suggest that SMEs differ with age (for reviews, see Friedman & Johnson, 2014; Maillet & Rajah, 2014a; Spreng, Wojtowicz, & Grady, 2010; Werkle-Bergner et al., 2006), adding weight to the proposal that age-related memory decline can be attributed, at least in part, to differences in how study events are processed at the time they are initially experienced.

The typical implementation of the subsequent memory procedure examines SMEs that reflect differences in the neural activity elicited by to-be-remembered study items. However, a now-substantial body of evidence from studies in healthy young adults indicates that successful encoding also depends on neural activity occurring in the second or so *before* a study item is encountered (Adcock, Thangavel, Whitfield-Gabrieli, Knutson, & Gabrieli, 2006; Addante, de Chastelaine, & Rugg, 2015; de Chastelaine & Rugg, 2015; Fernandez, Brewer, Zhao, Glover, & Gabrieli, 1999; Galli, Choy, & Otten, 2012; Galli, Gebert, & Otten, 2013; Gruber & Otten, 2010; Otten, Quayle, Akram, Ditewig, & Rugg, 2006; Otten, Quayle, & Puvaneswaran, 2010; Padovani, Koenig, Brandeis, & Perrig, 2011; Padovani, Koenig, Eckstein, & Perrig, 2013; Park & Rugg, 2010; for review, see N. Cohen et al., 2015). Crucially, it is currently unknown if, like SMEs, these ‘prestimulus’ SMEs (preSMEs) differ according to age. The present experiment addresses this question using ERPs to measure encoding-related neural activity.

### Prestimulus Subsequent Memory Effects

The typical ERP paradigm used to investigate preSMEs is best illustrated by a series of experiments reported by Otten and colleagues (2006; 2010). In these experiments participants were presented with a study task where each study item was preceded by a *task cue* that gave information about the upcoming study item, such as whether to make a semantic or an orthographic judgment about the item (Experiment 1 in Otten et al., 2006), or whether the item would be presented visually or auditorily (Experiment 2 in Otten et al., 2006; Otten et al., 2010). Prestimulus SMEs were examined by time-locking ERPs to the onset of the task cues and segregating the waveforms according to subsequent memory performance for the items following each cue. ERPs elicited by cues preceding later remembered items differed from those elicited by cues preceding forgotten items by virtue of a negative voltage shift over frontal midline electrodes maximal in the second or so prior to the onset of the study item.

Despite the strong evidence for preSMEs, their functional significance is not well understood. The most frequently discussed roles for the neural processing reflected by preSMEs are that the effects are either manifestations of spontaneous fluctuations in neural states that are differentially conducive to memory encoding (e.g., Ezzyat et al., 2017; Guderian, Schott, Richardson-Klavehn, & Duzel, 2009; Fernandez et al., 1999; Yoo et al., 2012; Fell et al., 2011) or the controlled engagement of task-specific preparatory processes that benefit memory encoding (e.g., de Chastelaine & Rugg, 2015; Galli et al., 2013; Otten et al., 2006; 2010). A full account of the functional role preSMEs play in memory encoding will likely incorporate aspects from both accounts. Findings from prior studies examining preSMEs associated with task cues (e.g., Otten et al., 2006), which is the approach taken in the present study, have provided support for the hypothesis that preSMEs reflect the engagement of preparatory processes. This view proposes that preSMEs reflect differential preparation of stimulus- or task-specific processes (the adoption of an appropriate ‘task set’) that facilitate formation of accessible memory representations (Addante et al., 2015; de Chastelaine & Rugg, 2015; Gruber & Otten, 2010;

Otten et al., 2006; 2010; Padovani et al., 2011; Park & Rugg, 2010). A critical finding that any hypothesis must account for is that preSMEs have consistently been reported for study items that were subsequently recollected, but not for items that were later recognized on the basis of an acontextual sense of familiarity in the absence of recollection (Addante et al., 2015; de Chastelaine & Rugg, 2015; Gruber & Otten, 2010; Otten et al., 2010; Padovani et al., 2011; Park & Rugg, 2010). That is, the current evidence suggests that preSMEs are best characterized as subsequent *recollection* effects.

The primary aim of the present study was to examine whether older adults demonstrate preSMEs associated with subsequent recollection like those that have been observed in young adults. Two lines of evidence favor the prediction that older adults should be impaired in engaging preparatory processes during encoding. First, as noted above, preSMEs appear to be selectively predictive of successful recollection (e.g., Park & Rugg, 2010), a form of memory highly vulnerable to aging (e.g., Koen & Yonelinas, 2016; for recent reviews, see Koen & Yonelinas, 2014; Schoemaker et al., 2014). Second, relative to young adults, older adults rely more heavily on ‘reactive’ rather than ‘proactive’ cognitive processing (e.g., Braver et al., 2001; Braver, Paxton, Locke, & Barch, 2009; Jacoby, Shimizu, Velanova, & Rhodes, 2005). That is, older adults tend to engage stimulus- or task-relevant cognitive processes only after a target event has been encountered. These lines of evidence lead us to predict that preSMEs will be reduced or absent in older adults.

### Post-Stimulus Subsequent Memory Effects

While the focus of the present study was on examining age differences in preSMEs, we also examined the effects of age on post-stimulus effects (SMEs). ERP studies employing the subsequent memory procedure have consistently reported positive SMEs onsetting between 300-600 ms post-stimulus for subsequently remembered versus forgotten study items (for reviews, see Friedman & Johnson, 2000; Rugg, 1995; Wilding & Ranganath, 2012). Whereas preSMEs typically show subsequent recollection effects, SMEs have been reported to be sensitive to both subsequent recollection and familiarity. At least in young adults, however, SMEs associated with recollection are typically larger than those associated with familiarity-based recognition (e.g., Cansino & Trejo-Morales, 2008; Duarte, Ranganath, Winward, Hayward, & Knight, 2004; Friedman & Trott, 2000; Mangels, Picton, & Craik, 2001).

As discussed previously, there are well-documented age differences in the magnitude of ERP SMEs (Friedman, Nessler, & Johnson, 2007; Friedman & Johnson, 2014; Werkle-Bergner et al., 2006). Although SMEs have been observed in older adults in association with simple item recognition (Gutchess, Ieuji, & Federmeier, 2007; Tellez-Alanis & Cansino, 2004), SMEs predictive of subsequent recollection (or high confidence recognition responses) are either reduced in magnitude (Friedman & Trott, 2000; Gutchess et al., 2007; Kamp & Zimmer, 2015), or differ in their onset and topography (Cansino, Trejo-Morales, & Hernandez-Ramos, 2010), in older adults relative to young adults. These findings have been taken as evidence that aging is associated with reduced efficacy of encoding processes that support later recollection (Craik & Rose, 2012; Friedman & Johnson, 2014; Werkle-Bergner et al., 2006). Here, we expected to replicate prior findings by finding an age-related attenuation in the magnitude of SMEs associated with subsequent recollection.

### The Present Experiment

The present study investigated age differences in preSMEs and SMEs using a procedure similar to that employed in Experiment 1 of Otten and colleagues (2006). Young and older adults incidentally encoded a list of words for a subsequent memory test. Each word in the study list was preceded by a task cue that instructed participants to make one of two possible semantic judgments about the word. At test, participants made a Remember/Know (Tulving, 1985) judgment for each word. Source memory for the encoding task was assessed for words attracting a ‘Remember’ judgment. Although prior research has typically used Remember/Know judgments to investigate ERP preSMEs (Otten et al., 2006; 2010), for reasons discussed in the Methods, we operationalized recollection as accurate versus inaccurate source memory for the encoding task. Prestimulus SMEs were assessed with ERPs time-locked to the onset of the task cue and segregated by whether the word was associated with accurate versus inaccurate source memory. SMEs were examined in a similar fashion with ERPs that were time-locked to the onset of the study words.

As discussed above, our prediction was that we would observe age differences in preSMEs. Specifically, we expected that preSMEs would be attenuated or undetectable in older adults. A failure to detect preSMEs in older adults could arise for several reasons, however, not all of which are of equal theoretical interest. For instance, the absence of prestimulus effects might indeed suggest age differences in the efficacy of preparatory processes, but alternately it might simply indicate that older adults did not attempt to engage these processes (Duverne, Motamedinia, & Rugg, 2009; Luo & Craik, 2008). A similar issue has arisen in research examining the engagement of ‘material-dependent’ retrieval orientations (Robb & Rugg, 2002) in young and older adults. Although initial results suggested that older adults do not engage specific retrieval orientations (Jacoby et al., 2005; Morcom & Rugg, 2004), subsequent research demonstrated that older adults are able to do so when suitably incentivized (Duverne et al., 2009). Considering these findings, in the present experiment we manipulated the presentation duration of the study items (300 ms vs. 1000 ms) in an effort to vary the incentive for participants to engage prestimulus preparatory processes. The assumption underlying this manipulation is that the relatively limited perceptual availability of the study words in the short duration condition, which is similar to the study durations employed in prior ERP studies examining preSMEs (e.g., Gruber & Otten, 2010; Otten et al., 2006; Otten et al., 2010), will place a premium on engaging preparatory processes in the prestimulus interval. Thus, our expectation was that the highest likelihood of finding preSMEs in older adults would be in the short encoding condition, when the incentive to engage prestimulus preparatory processes is greatest because participants need to devote more resources to accurately perceiving and processing the word. We note however that under the hypothesis that preSMEs reflect spontaneous neural states or fluctuations that benefit memory encoding processes (see above), this manipulation would be predicted to have a null effect on the magnitude of preSMEs.

## Materials and Methods

### Ethics Statement

The Institutional Review Board of the University of Texas at Dallas approved the experimental procedures described below. All participants provided written informed consent prior to participating in each experimental session.

### Participants

Twenty-four older and 24 young participants were included in the analyses reported in this paper (see Table 1 for characteristics of the sample). Participants were recruited from the University of Texas at Dallas and the greater Dallas metropolitan area and compensated for travel expenses and time (at the rate of $30/hour). All participants were right-handed, reported having normal or corrected-to-normal vision, and scored 27 or more on the Mini-Mental State Examination (MMSE; Folstein, Folstein, & McHugh, 1975). Exclusion criteria included a history of cardiovascular disease (other than treated hypertension), psychiatric disorder, illness or trauma affecting the central nervous system, substance abuse, and self-reported current or recent use of psychotropic medication or sleeping aids.

**Table 1.** Demographic and neuropsychological test data for young and older adults.

|  | Young Adults | Older Adults | *p*-value |
| --- | --- | --- | --- |
| Sex (M/F) | 11/13 | 11/13 | - |
| Age | 23.50 (3.27) | 69.42 (3.83) | - |
| Education | 16.25 (2.03) | 17.33 (2.96) | 0.147 |
| MMSE | 29.54 (0.66) | 29.21 (0.83) | 0.131 |
| CVLT Short Delay – Free | 13.17 (1.76) | 10.75 (3.38) | 0.004 |
| CVLT Short Delay – Cued | 13.75 (1.70) | 11.83 (2.78) | 0.006 |
| CVLT Long Delay – Free | 13.83 (1.76) | 11.13 (3.33) | 0.001 |
| CVLT Long Delay – Cued | 14.17 (1.49) | 11.88 (3.00) | 0.002 |
| CVLT Recognition – Hits | 15.50 (0.78) | 14.88 (1.42) | 0.067 |
| CVLT Recognition – False Alarms | 0.54 (0.66) | 2.13 (2.35) | 0.004 |
| Logical Memory I | 30.29 (4.32) | 28.63 (6.50) | 0.302 |
| Logical Memory II | 27.71 (5.20) | 25.08 (6.75) | 0.139 |
| Digit Span Total<sup>1</sup> | 21.54 (3.82) | 18.96 (3.18) | 0.014 |
| Symbol-Digit | 61.17 (12.67) | 51.04 (7.97) | 0.002 |
| Trails A (secs) | 21.96 (8.64) | 26.93 (8.68) | 0.053 |
| Trails B (secs) | 52.07 (21.46) | 65.22 (21.58) | 0.040 |
| FAS Total | 44.54 (9.35) | 45.54 (8.33) | 0.697 |
| Category Fluency (Animals) | 22.71 (4.91) | 20.71 (5.84) | 0.206 |
| WTAR (Raw) | 40.58 (3.24) | 43.46 (4.12) | 0.010 |
| Raven's (List 1) | 11.00 (1.06) | 8.83 (2.04) | < 0.001 |

*Note.* Standard deviations are reported in parentheses. The *p*-values were obtained from Welch *t*-
tests comparing young and older adults. <sup>1</sup>Digit span total equals the sum of forward and backward span.

Data from an additional 7 participants who underwent EEG recording were excluded for the following reasons. Specifically, 2 young adult females and 2 older adult males were excluded due to insufficient (< 15) artifact-free epochs in a bin of interest, 1 young adult male and 1 older adult male were excluded because of significant artifacts in the ERP that could not be corrected or removed (as jointly determined by the first and last author), and 1 older adult male was excluded for incorrect use of Remember/Know judgments.

### Neuropsychological Testing

Participants completed a neuropsychological test battery on a separate day prior to the EEG study. The battery included the MMSE, California Verbal Learning Test-II (CVLT; Delis, Kramer, Kaplan, & Ober, 2000), the digit symbol, and forward and backward digit span subtests of the Wechsler Adult Intelligence Scale – Revised (Wechsler, 1981), trail making tests A and B (Reitan & Wolfson, 1985), the F-A-S subtest of the Neurosensory Center Comprehensive Evaluation for Aphasia (Spreen & Benton, 1977), the category fluency test for animals (Benton, 1968), Wechsler test of adult reading (WTAR; Wechsler, 2001), the logical memory subtest of the Wechsler Memory Scale (Wechsler, 2009), and List 1 of the Raven’s Progressive Matrices (Raven, Raven, & Courth, 2000). Volunteers were excluded from participating in the EEG study if (1) one or more of the memory measures (i.e., CVLT or logical memory) were more than 1.5 standard deviations below the age- and education-adjusted mean, (2) they had a standard score below 100 on the WTAR, or (3) two or more scores on non-memory tests were 1.5 standard deviations below the mean. Additionally, participants completed a visual acuity test as part of the battery. However, this measure was not used for screening purposes and is not relevant to the current study, and thus it will not be discussed further.

### Materials

Three hundred and eighty-four nouns from the MRC Psycholinguistic database (Coltheart, 2007) comprised the critical stimuli in this experiment. The words ranged from 4-8 letters in length (*M* = 5.32, *SD* = 1.21), from 1-40 in Kucera-Francis frequency (*M* = 13.33, *SD* = 10.25) (Kucera & Francis, 1967) and concreteness ratings from 500-662 (*M* = 583.90, *SD* = 31.99). These words were used to generate 24 stimulus sets that were yoked across young and older participants.

For each stimulus set, 256 words were assigned to be presented in the study list and were randomly divided into 4 groups of 64 words. The four groups were assigned to one of the four cells formed by crossing the encoding duration variable (short versus long encoding duration) and the semantic judgment performed during the study phase (Manmade versus Shoebox). The word list for the test phase comprised the 256 words from the study phase and remaining 128 words as new items on the recognition memory test.

An additional 24 words with similar characteristics to those of the critical stimuli were used in the practice phases. The words in each practice list were the same for all participants. In total there were 3 practice study lists (self-paced, speeded, real), each comprising 8 words. In addition, there were two 2 practice test lists (feedback, real). The feedback practice test list consisted of 8 words from the practice study phases and 4 new words. The real practice test phase consisted of 8 words from the practice study phases and 4 new words.

Stimuli were presented via a 22” LCD Monitor (1024 x 768 resolution) using Cogent 2000 software (www.vislab.ucl.ac.uk/cogent_2000.php) implemented in Matlab 2012b (www.mathworks.com). Throughout the experiment, the monitor displayed a black background with a white centrally located box (250 x 250 pixels). All stimuli were presented in the center of the screen in Helvetica 32-point font. Note that, as is typical in studies such as this, all timing information given below is uncorrected for the refresh rate of the monitor, and hence varied across trials within a range of ±17ms.

### Procedure

#### Overview

The experiment was completed across two sessions on different days, with the neuropsychological test battery completed in the first session, and the experimental EEG session completed in the second session. The EEG memory task comprised study and test phases structured similarly to previous studies (Galli et al., 2013; Otten et al., 2006; 2010).

#### Study Phase

The procedure for the critical study phase is schematically depicted in the left half of Figure 1. Participants were presented with four blocks of 64 words that served as the studied items for the subsequent memory test. Half of the blocks were assigned to the short encoding condition, and the remaining half were assigned to the long encoding condition. A break was given halfway through each block. The instructions did not reference the subsequent memory test; therefore encoding was incidental. Each word in both the short and long encoding conditions was preceded by a task cue informing participants which one of two judgments they should make about the word. The two possible judgments were ‘Manmade’, whereby participants made a yes/no response as to whether the object denoted by the word was a manmade object, and ‘Shoebox’, which required a yes/no response about whether the object denoted by the word was something that fits inside a shoebox. The cues for the Manmade and Shoebox tasks were a red ‘X’ and ‘O’, respectively. In both the short and long blocks, each trial began with a green fixation cross for 500 ms, which was followed by the presentation of the task cue for 500 ms. After a 1500 ms delay (filled with a black fixation cross), the study word was presented. In the short encoding condition, the word appeared on the screen for 300 ms followed by a 2700 ms black fixation cross that served as the response window. In the long encoding condition, the word appeared on the screen for 1000 ms followed by a 2000 ms black fixation cross that served as the response window. The hand that was used for the Manmade as opposed to the Shoebox judgment was counterbalanced across participants, and they used their index or middle fingers to enter a yes or no response, respectively. Note that the timing of the study trials and inter-trial interval was fixed and hence unaffected by response latency. Participants were instructed to make their responses as quickly but as accurately as possible.

**Figure 1.**
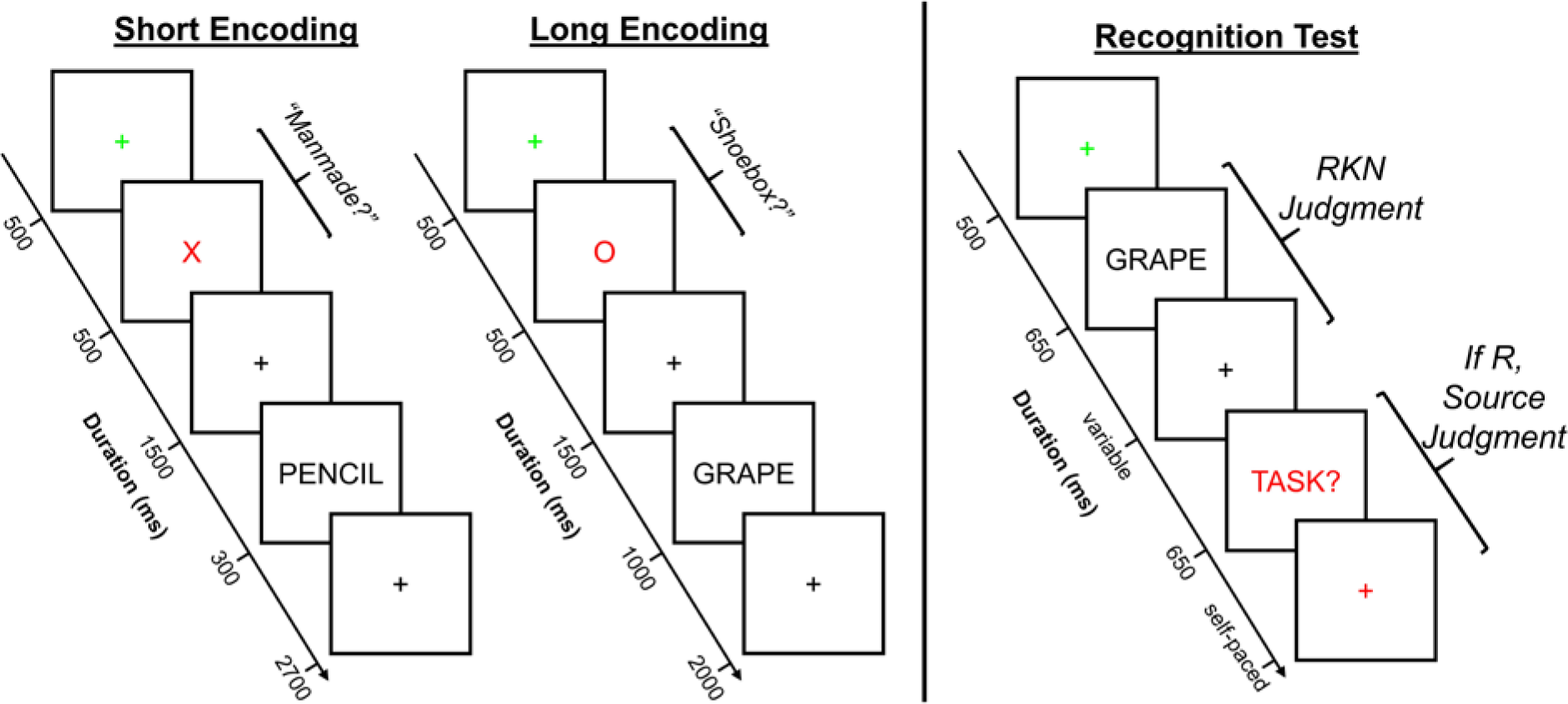
Visual depiction of the experimental memory task. During the study phase, participants were presented with a list of words and asked to make one of two possible judgments about each word. Prior to the presentation of each word, participants were presented with a task cue, specifically a red ‘X’ or ‘O’, that instructed participants to make either the Manmade or Shoebox judgment, respectively, about the following word. There were two blocked encoding conditions (short and long) that varied in the amount of time the word appeared on the screen, and participants were informed in advance of the duration of the word. Importantly, the amount of time participants had to respond did not differ between the two encoding conditions. At test, participants were shown words from the study phase intermixed with new words. Participants first made an old/new decision using Remember, Know, and New judgments and, when a Remember response was given, participants were asked to retrieve the judgment they made about the word during the study phase. The response alternatives for this latter judgment included the label of the two encoding tasks (Manmade and Shoebox) as well as a Don’t Know response option. Participants were not informed about the subsequent test prior to the study phase, and thus encoding was incidental. EEG was recorded during both the study and test phases.

Participants completed three brief practice study phases prior to the critical phase. The three practice study phases comprised (in order) a self-paced practice with feedback, speeded practice with feedback, and a “real” practice phase. In the self-paced practice phase, participants were presented with the trial sequence as described above with the exception that the word remained on the screen until a response was given. After entering a response, participants received feedback as to whether they responded to the correct judgment (i.e., they entered their judgment using the assigned hand for the Manmade or Shoebox judgments). Feedback as to the accuracy of the yes and no response for either judgment was not provided. In the speeded practice with feedback phase, participants were presented with the full trial sequence described above for the critical study phase, with the exception that after each trial participants were provided feedback as to whether they responded to the correct judgment. In this practice phase, participants were also introduced to the short and long encoding conditions. The purpose of the first two practice study phases was to (1) practice participants in using different hands to answer the two different judgments and (2) instruct participants to respond quickly. The “real” practice phase was identical to the critical study phase in timing, except that the blocks were shorter (4 trials in each encoding condition).

#### Test Phase

The procedure for the test phase is depicted in the right half of Figure 1. The test phase instructions began approximately 15 minutes after the study phase. In the test phase proper (see below for description of the practice phases) participants were presented with studied and new words one at a time and required to make a Remember/Know/New (R/K/N) judgment (Tulving, 1985). Participants were instructed to make a Remember (R) response if they could recollect at least one specific detail from the study phase that was associated with the test item (e.g., the encoding task; a thought that came to mind). Importantly, participants were further instructed that they should be able to explain exactly what they recollected about the word to the experimenter if they made an R response. A Know (K) response was required when participants had high confidence that the word was studied but they were unable to recollect any specific contextual detail about the word when it was encountered in the study phase. The K response was labeled as ‘Familiar’ in the instructions to participants to avoid confusion between the terms Remember and Know, but we refer to this response option here as Know (or K) to remain consistent with the literature. Participants were instructed to give a New (N) response if they believed the word did not appear in the list or if they would simply be guessing that the word had been studied.

For words endorsed with an R response, participants were further required to make a source memory judgment in respect of the encoding task (Manmade or Shoebox). This judgment had three possible responses: Manmade, Shoebox, and Don’t Know. Participants were instructed to select Manmade or Shoebox if they had high confidence about the task they had performed on the word during encoding. They were instructed to use the Don’t Know option if they were not confident or were unable to recollect the encoding task. Thus, an R response followed by a Don’t Know source memory response indicated that participants recollected a specific detail or details about the study that did not include the criterial details necessary to support a high confidence source memory decision (i.e., the nature of the encoding task).

It should be noted that the present task differs from prior studies using combined R/K/N and source memory judgments (e.g., Duarte, Henson, & Graham, 2008) in two ways. First, we included a Don’t Know response alternative on the source memory judgment. Second, source memory judgments were required for trials receiving an R judgment, but not for trials receiving a K judgment. We adopted this procedure on the assumption that, as participants were instructed to give a K response if they were unable to recollect or retrieve *any* specific detail of the study episode, the great majority of source judgments following a K response would have led to a Don’t Know response. By restricting the source judgments to items accorded an R judgment we could markedly simplify the task instructions and response demands, reducing the likelihood of non-compliance or confusion.

Each trial in the test phase began with a green fixation cross for 500 ms, which was followed by the presentation of the test word for 650 ms, and lastly a black fixation cross. Participants were instructed to enter their R/K/N response after the black fixation cross appeared on the screen. The R/K/N response was self-paced, and thus the fixation cross remained on the screen until a response was entered. If participants made a K or N response, the fixation cross remained for an additional 2000 ms, after which the next trial began (i.e., the next green fixation cross). If participants made an R response, the black fixation cross remained on the screen for an additional 500 ms, and was followed by a prompt (the word ‘Task?’ in red font) to enter the source memory response. The prompt remained on the screen for a maximum of 650 ms, and was replaced with a red fixation cross if a response was still forthcoming. The source memory response was also self-paced. Once the response was entered, a black fixation cross appeared for 1000 ms before the onset of the next trial. The test phase was broken into 12 equal-length blocks, allowing a short break after every 32 trials.

Prior to the critical test phase there were two practice phases that made use of the words from the practice study phases intermixed with a set of new words. The timing for both practice phases followed that described in the preceding paragraph for the critical phase. In the first practice phase, the participant’s response selections were displayed on the computer screen at the end of each trial, and they were asked to explain the basis of their judgments. This was to allow the experimenters to provide participants with feedback on their use of the R/K/N responses, and to verify that participants were applying the R/K/N distinction correctly. The second practice phase was identical to the test phase proper, except that a shorter list was employed. Following completion of the critical test phase, participants were debriefed, compensated, and thanked for their participation.

### Rationale for the Test Phase

The rationale for including both R/K/N and source memory judgments was to explore whether preSMEs and SMEs differed for R judgments associated with accurate versus inaccurate source judgments. That is, we intended to contrast accurate source memory controlling for subjective reports of recollection (i.e., compare R judgments with accurate and inaccurate source memory), as well as to contrast subjective reports of recollection accompanied by inaccurate source memory with familiarity-based recognition (i.e., K responses). Contrary to our expectations based on the results from a behavioral pilot study, limited trial counts precluded us from being able to address this question. Instead, we examined pre- and post-stimulus SMEs by operationalizing successful versus unsuccessful recollection in terms of correct and incorrect source memory judgments, respectively. Specifically, source correct trials comprised encoding trials for which the study word went on to receive both an R response and a correct source memory decision. Source incorrect trials comprised trials where the study word subsequently received an R response and either an incorrect or a Don’t Know response to the source memory judgment, or trials that subsequently received a K response. Given our assumption that participants would have responded Don’t Know (or guessed) on trials accorded a K judgment, this partitioning of trials is analogous to that employed in prior studies where source memory judgments were not combined with the R/K/N procedure (e.g., Koen & Rugg, 2016; Maillet & Rajah, 2014b; Mattson, Wang, de Chastelaine, & Rugg, 2014; Ranganath et al., 2004).

### EEG/ERP Recording and Analysis

EEG was recorded continuously during both the study and test phases (only the study data are presented here). The recordings were made from a total of 64 Ag/Ag-Cl electrodes with 58 embedded in an elastic cap (EasyCap; Herrsching-Breitbrunn, Germany; www.easycap.de; montage11) and 6 affixed directly to the skin. The electrode sites in the cap comprised 6 midline locations (Fpz, Fz, Cz, CPz, PZ, POz) and 26 homotopic electrode pairs (Fp1/2, AF3/4, AF7/8, F1/2, F3/4, F5/6, F7/8, FC1/2, FC3/4, FC5/7, FT7/8, C1/2, C3/4, C5/6, T7/8, CP1/2, CP3/4, CP5/6, TP7/8, P1/2, P3/4, P5/6, P7/8, PO3/4, PO7/8, and O1/O2). Two additional electrodes were affixed to the left and right mastoid process. Vertical and horizontal EOG were monitored with electrode pairs above and below the right eye, and on the outer canthi of the left and right eyes, respectively. The ground and online reference electrodes were embedded in the cap at sites AFz and FCz, respectively. EEG and EOG channels were digitized at 500 Hz using an amplifier bandpass of .01-70 Hz (3dB points) using the BrainVision Recorder software package (version 1.20.0601; www.brainvision.com). Electrode impedances were adjusted to be less than or equal to 5 kΩ prior to the start of the study phase, and were rechecked and, if necessary, adjusted, during each break.

The EEG data were processed offline in Matlab 2012b (www.mathworks.com) using EEGLAB version 13.5.4 (Delorme & Makeig, 2004), ERPLAB version 5.0.0.0 (Lopez-Calderon & Luck, 2014), and custom Matlab code. The continuous EEG data were digitally filtered between .03 and 19.4 Hz with a zero-phase shift butterworth filter (12 dB/octave roll-off, DC offset removed prior to filtering) using ERPLAB. Pre- and post-stimulus EEG epochs were processed separately, albeit in identical processing streams. Epochs were extracted with a total duration of 2500 ms (from −500 ms to 2000 ms relative to the onset of the task cue or the study word). The epoched data were subjected to Independent Components Analysis (ICA; Jung et al., 2000) to identify artifactual EEG components (e.g., blinks, eye movements, muscle). Prior to ICA, the epochs were baseline corrected to the average voltage across the epoch to improve estimation of the ICA components (Groppe, Makeig, & Kutas, 2009), and epochs with non-stereotypical artifacts (e.g., due to a participant coughing) were rejected. Rejection of the entire data belonging to an electrode, when necessary, was conducted prior to ICA. Data from rejected electrodes were replaced using Spline interpolation after removal of artifactual ICA components. The SASICA (Chaumon, Bishop, & Busch, 2015) and ADJUST (Mognon, Jovicich, Bruzzone, & Buiatti, 2011) EEGLAB plug-ins were used to aid with the identification of artifactual components, and the final determination as to whether to remove a component was made by the first author. After ICA artifact correction, the epoched EEG data were re-referenced to linked mastoids (recovering the FCz electrode) and baseline corrected to the average voltage of the 500 ms preceding the time-locked event (the task cue or the study word). Epochs were rejected from averaging if (1) voltage in the epoch exceeded ±100 μV, (2) baseline drift exceeded 40 μV (determined as the absolute difference in amplitude between the average amplitude of the first and last 250 ms of each epoch), (3) an artifact was present based on visual inspection by the first author, (4) participants failed to respond to or incorrectly responded (i.e., used the incorrect hand) on the study judgment, or (5) a participant’s reaction time (RT) was faster than 450 ms during the study phase or 650 ms during the test phase.

ERPs for each electrode site and event of interest (task cue and study word) were created by averaging all artifact-free epochs according to the subsequent memory judgment within each encoding condition (short vs. long). For the reasons discussed above (see Rationale for the Test Phase), we segregated trials into three subsequent memory bins: (1) trials receiving an R response in conjunction with a correct source judgment, (2) trials receiving an R response in conjunction with an incorrect or Don’t Know source memory decision, along with trials receiving a subsequent K response, and (3) trials receiving a subsequent New response (i.e., forgotten, or inaccessible, study words). The above binning scheme was applied separately to the short and long encoding conditions, resulting in a total of six bins for each event of interest (task cue and study word). For several of the participants (7 young and 6 older), fewer than 15 artifact-free trials were available for the forgotten condition. Therefore, this bin was excluded from the group analysis and will not be discussed further.

### Statistical Analyses

Statistical analyses were carried our using *R* software (version 3.4.3; R Core Team, 2015). ANOVAs were conducted using the *afex* package (version 0.16-1; Singmann, Bolker, Westfall, & Aust, 2016). The Greenhouse-Geisser procedure was used to correct the degrees of freedom for non-sphericity in the ANOVAs (Greenhouse & Geisser, 1959), and is reflected in the reported degrees of freedom. Post-hoc tests on significant interactions involving Memory were conducted using the *lsmeans* package (version 2.27.61; Lenth, 2016) with degrees of freedom estimated using Satterthwaite (1946) approximation. Effect size measures for results from the ANOVAs are reported as partial-η^2^ (J. D. Cohen, 1988). Results were considered significant at *p* < .05.

#### Behavioral Data

The dependent variables of interest from the behavioral data included three estimates of memory performance derived from the test data, and the median reaction time (RT; in ms) to the judgment during the study phase. RT for each participant was computed as the median RT for trials from the four bins of interest in the ERP analysis.

The three memory estimates included estimates of recollection and familiarity derived from the R/K/N responses and source memory accuracy derived from the source memory judgments following an R response. Estimates of recollection and familiarity were calculated (without regard to source memory accuracy) separately for each encoding condition using the independent R/K/N estimation procedure (Yonelinas & Jacoby, 1995). Recollection was computed with the following formula:

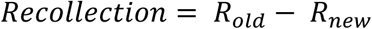

*R_old_* and *R_new_* represent the proportion of R responses to old and new items, respectively. Familiarity estimates were derived from the following formulae:

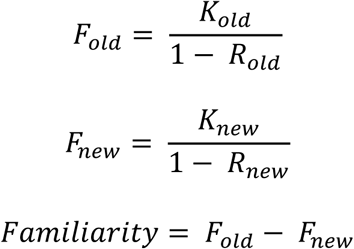

*K_old_* and *K_new_* represent the proportion of K responses to old and new items, respectively.

Source memory was computed using a single-high threshold model (Snodgrass & Corwin, 1988) modified to account for the “guess” rate (e.g., Mattson et al., 2014). Source accuracy was computed separately for the short and long conditions as follows:

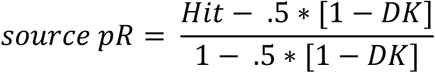

Two different measures of source pR were calculated. First, we calculated pR restricted to items receiving an R response, such that the *Hit* and *DK* variables in the above formula refer to the proportion of R responses accompanied by an accurate or don’t know source memory, respectively. This measure will hereafter be referred to as pR_Rem_. Additionally, for consistency with the binning scheme of the ERP analysis, we computed a pR measure whereby the *Hit* and *DK* proportions were computed conditional on giving an R or K response (i.e., an ‘old’ response) to studied words. This measure, which we hereafter refer to as pR_ERP_, treats K responses as don’t know source judgments based on our assumption that participants would have given a don’t know response if asked to do so (see Rationale for the Test Phase).

#### ERP Analysis

The analysis of the ERP data focused on preSMEs (time-locked to the onset of the task cues) and SMEs (time-locked to the onset of the study words) in specific time windows, which will be discussed further below. In all time windows, ERP amplitude was computed as the mean voltage (in μV) in the time window of interest relative to the mean voltage in the 500 ms time window prior to the time-locked event (task cue or study word). These raw mean voltages were submitted to the ANOVAs unless otherwise specified.

As discussed in the Introduction, preSMEs in young adults typically onset approximately one second after presentation of the task cue and increase in magnitude until the presentation of the study item. However, prior studies have found some variability in the time window showing the maximal preSME with reports that preSMEs are maximal immediately before onset of the study item (e.g., Otten et al., 2006) or that the effect is maximal between 500 and 1000 ms prior to onset of the study item (e.g., Galli et al., 2013; Otten et al., 2010). To avoid circularity with selecting the time window based on observing the current data, we *a priori* defined three equal length time windows (500-1000 ms, 1000-1500 ms, and 1500-2000 ms) to examine preSMEs.

We expected preSMEs would be most prominent in the latter two windows as these correspond to the one second interval before the onset of the study event. The earlier time window was included as a control with the expectation that no reliable preSMEs would be present in either age group.

Prior research has established positive-going ERP SMEs are typically observed for trials subsequently recollected compared to those recognized in the absence of recollection (e.g., Cansino & Trejo-Morales, 2008; Cansino et al., 2010; Friedman & Trott, 2000). This positive going effect typically onsets approximately 600 ms after the study item, and is sustained throughout much of the epoch. However, some studies have shown SMEs to onset as early as 300 ms following presentation of a study item. Therefore, three *a priori* time windows (300-600 ms, 600-1500 ms, and 1500-2000 ms) typical of prior studies were used to examine the SMEs.

The data from each of the time windows described above were submitted to separate omnibus ANOVAs that included the between participant factor of Age (young, old), and the within participant factors of Memory (source correct, source incorrect), and Encoding Condition (short, long), along with three within-subject electrode site factors: anterior/posterior (AP) chain (Fp, F, C, P, O), laterality (medial, lateral), and hemisphere (left, right). The electrodes included in the analysis, and how they were assigned to the three electrode site factors, are depicted in Figure 2. Given our interest in examining subsequent memory effects, only significant effects from the omnibus ANOVAs including the Memory factor are reported.

**Figure 2.**
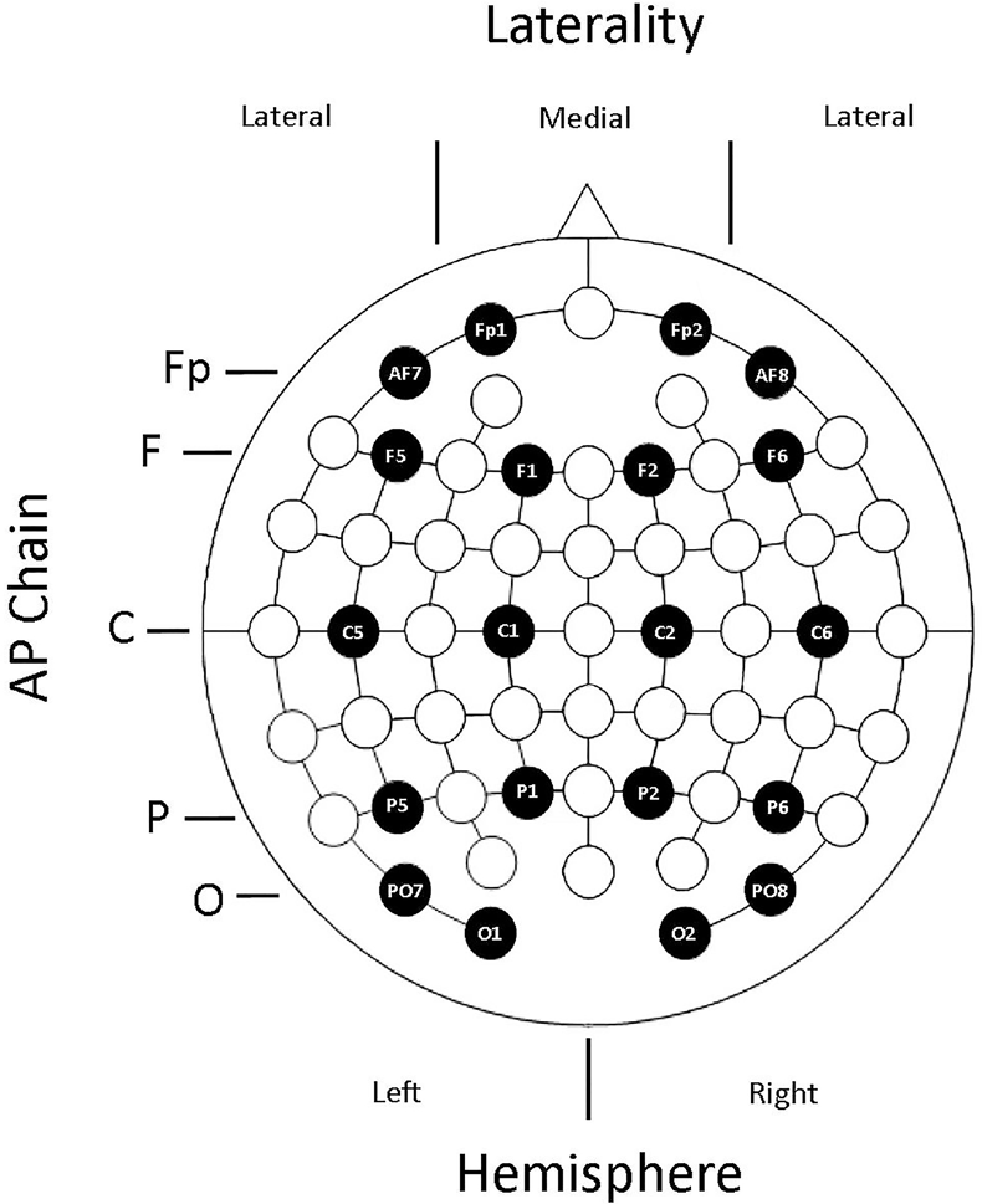
Visual depiction of the electrodes and the anterior-posterior chain (AP Chain), Laterality, and Hemisphere electrode factors used in the ERP omnibus ANOVAs.

To foreshadow the results, significant effects from the omnibus ANOVAs either included interactions between Memory and Age, or between Memory, Age, and Encoding condition. Significant interactions involving Memory and Age were followed up with separate ANOVAs in each age group that included the factors Memory, AP Chain, Laterality, and Hemisphere (after collapsing the data across the two encoding conditions).

Interactions from the omnibus ANOVA involving Age, Memory, and Encoding condition were first followed up with separate Memory, Age, AP Chain, Laterality, and Hemisphere ANOVAs for each encoding condition. The ANOVAs were split across the Encoding factor rather than Age because our primary focus was on examining age differences in both preSMEs and SMEs. Interactions in a given encoding condition (short or long) in the above-described ANOVAs involving Age and Memory were further decomposed with separate Memory, AP Chain, Laterality, and Hemisphere ANOVAs for the young and older samples.

Lastly, we aimed to establish simple effects of memory (both pre- and post-stimulus) in time windows showing significant effects involving the Memory factor in the omnibus ANOVA. For preSMEs, which have an anterior frontal distribution (e.g., Otten et al., 2006), simple effects of memory were evaluated by conducting contrasts on the least-squares means (or estimated marginal means) over electrodes Fp1, Fp2, F1, and F2, unless otherwise specified. Post-stimulus SMEs have fronto-central distribution (Cansino & Trejo-Morales, 2008; Cansino et al., 2010; Friedman & Trott, 2000), and therefore simple effects of memory were assessed over electrodes F1, F2, C1, and C2. Note that these contrasts were conducted using pooled variance measures in the context of the ANOVA model that produced the significant effect involving Memory.

## Results

### Neuropsychological Test Performance

The results from the various measures of the neuropsychological test battery are reported in Table 1. As in prior studies from our lab (de Chastelaine, Mattson, Wang, Donley, & Rugg, 2016; Mattson et al., 2014; Wang, de Chastelaine, Minton, & Rugg, 2012), older adults performed significantly worse than the young adult sample on tests assessing declarative memory, reasoning ability, and processing speed, but were equally proficient at word reading, as well as verbal and category fluency.

### Behavioral Performance

#### Study RT

Table 2 shows the group averages of each participant's median reaction time (RT) during the study phase as a function of subsequent memory. The ANOVA on the median RTs did not reveal any effects that included Age, all *p’s* involving Age ≥ 0.177. The absence of age effects suggests that any age differences in SMEs are unlikely to be attributable to group differences in RT. The ANOVA did however reveal two significant main effects. There was a main effect of Memory, F(1,46) = 6.40, *MSe* = 6847, *p* = 0.015, partial-η^2^ = 0.12, with slower RTs on trials in the source correct bin (*M* = 1247 ms, *SE* = 34 ms) relative to trials in the source incorrect bin (*M* = 1217 ms, *SE* = 33 ms). Additionally, there was a main effect of Encoding condition, F(1,46) =6.06, *MSe* = 14592, *p* = 0.018, partial-η^2^ = 0.12, that was driven by faster RTs for study trials belonging to the long (*M* = 1210 ms, *SE* = 33 ms) than the short (*M* = 1253 ms, *SE* = 34 ms) condition. It is important to point out that the slower RTs in the short encoding condition are consistent with the assumption presented in the Introduction that limited perceptual availability increases task difficulty, and therefore might increase incentive for the engagement of prestimulus processes.

**Table 2.** Mean (and standard errors) for the average median reaction time (RT) to judgments made during the study phase.

|  | Young Adults |  | Older Adults |  |
| --- | --- | --- | --- | --- |
|  | Short | Long | Short | Long |
| Source Correct | 1257 (74) | 1185 (73) | 1289 (69) | 1229 (63) |
| Source Incorrect | 1237 (72) | 1177 (73) | 1256 (59) | 1223 (60) |

Note. The bins for the RTs were comprised of the same trials as the ERP bins.

#### Test Performance

Table 3 lists the proportions of test trials receiving R (separated by the accuracy of the source memory response), K, and N responses. The numerical trend is that, relative to young adults, older adults showed (1) a decrease in the overall proportion of R responses to studied items, (2) a decreased rate of R responses that attracted a correct (R+SC) or a “don’t know” source memory decision (R+DK), and (3) an elevated rate of R responses associated with an incorrect source memory decision (R+SI). Although the overall rate of incorrect R responses to new items was relatively consistent across the two age groups, older adults were more likely to endorse a new item as coming from one of the encoding tasks relative to young adults. In addition to these trends, there were minimal differences between the two age groups in the raw proportion of K responses given to old and new items.

**Table 3.**
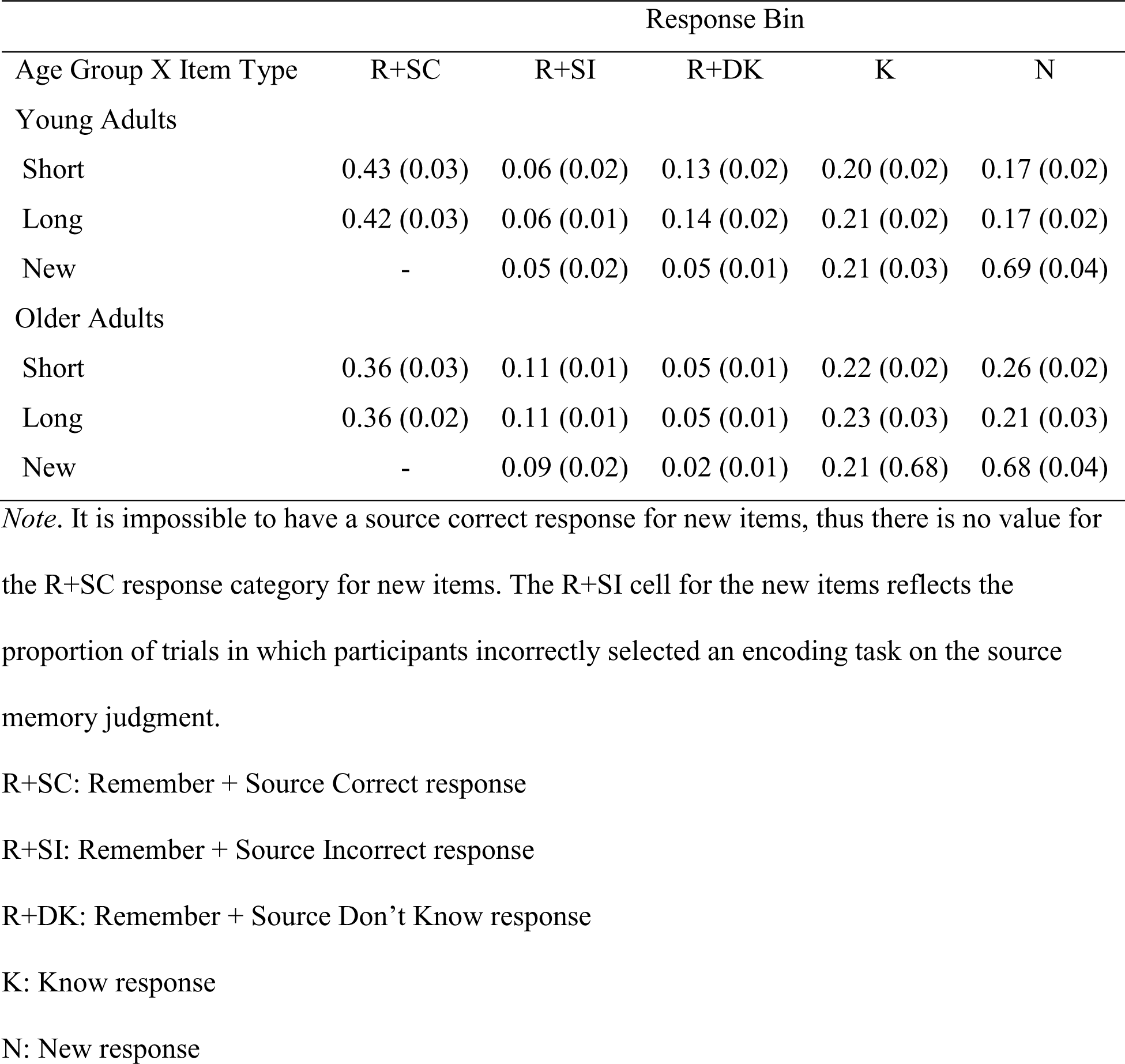
Means (and standard errors) for the proportion of trials in each response category.

| Age Group X Item Type | Response Bin |  |  |  |  |
| --- | --- | --- | --- | --- | --- |
|  | R+SC | R+SI | R+DK | K | N |
| Young Adults |  |  |  |  |  |
| Short | 0.43 (0.03) | 0.06 (0.02) | 0.13 (0.02) | 0.20 (0.02) | 0.17 (0.02) |
| Long | 0.42 (0.03) | 0.06 (0.01) | 0.14 (0.02) | 0.21 (0.02) | 0.17 (0.02) |
| New | - | 0.05 (0.02) | 0.05 (0.01) | 0.21 (0.03) | 0.69 (0.04) |
| Older Adults |  |  |  |  |  |
| Short | 0.36 (0.03) | 0.11 (0.01) | 0.05 (0.01) | 0.22 (0.02) | 0.26 (0.02) |
| Long | 0.36 (0.02) | 0.11 (0.01) | 0.05 (0.01) | 0.23 (0.03) | 0.21 (0.03) |
| New | - | 0.09 (0.02) | 0.02 (0.01) | 0.21 (0.68) | 0.68 (0.04) |

*Note.* It is impossible to have a source correct response for new items, thus there is no value for the R+SC response category for new items. The R+SI cell for the new items reflects the proportion of trials in which participants incorrectly selected an encoding task on the source memory judgment.
R+SC: Remember + Source Correct response
R+SI: Remember + Source Incorrect response
R+DK: Remember + Source Don't Know response
K: Know response
N: New response

Relevant to the ERP analyses reported below, age group differences in the relative proportion of trials contributing to the source incorrect bin would confound any group differences in the ERPs. This could occur, for example, if the relative proportion of trials comprising the source incorrect bin differed between the two age groups. Although there is no apparent difference in the proportion of K trials contributing to the source incorrect bin, older adults were less likely to provide a Don’t Know response for the source memory judgment following an accurate R response (i.e., fewer R+DK trials). This pattern of responding is quite common in source memory tasks that employ a DK response option (e.g., Dulas, Newsome, & Duarte, 2011; Dulas & Duarte, 2012; Mattson et al., 2014), and it is conceivable that it introduces a confound when comparing ERPs obtained from young and older adults. This issue should be kept in mind when interpreting the ERP results reported below.

Prior research has typically demonstrated large age differences in recollection estimates accompanied by smaller, albeit significant age differences in familiarity estimates derived from R/K/N judgments (Koen & Yonelinas, 2014). This pattern of results was observed here (Table 4). Specifically, there were significant age-related reductions in both recollection, F(1,46) = 9.00, *MSe* = 0.04, *p* = 0.004, partial-η^2^ = 0.16, and familiarity, F(1,46) = 4.36, *MSe* = 0.05, *p* = 0.042, partial-η^2^ = 0.09. There were no significant effects or interactions involving the Encoding factor on recollection or familiarity, all p’s ≥ 0.240.

**Table 4.** Means (and standard errors) for the mnemonic process estimates derived from the Remember/Know/New (R/K/N) and source memory judgments.

| Age Group X Item Type | Recollection | Familiarity | pR<sub>Rem</sub> | pR<sub>ERP</sub> |
| --- | --- | --- | --- | --- |
| Young Adults |  |  |  |  |
| Short | .053 (0.03) | 0.30 (0.04) | 0.51 (0.04) | 0.32 (0.03) |
| Long | 0.52 (0.03) | 0.31 (0.03) | 0.48 (0.05) | 0.31 (0.03) |
| Older Adults |  |  |  |  |
| Short | 0.40 (0.03) | 0.20 (0.03) | 0.45 (0.05) | 0.27 (0.03) |
| Long | 0.40 (0.03) | 0.22 (0.03) | 0.47 (0.04) | 0.26 (0.03) |

*Note.* The columns for Recollection and Familiarity refer to the estimates derived from the R/K/N judgments. The pR<sub>Rem</sub> and pR<sub>ERP</sub> were computed from the source memory judgments conditionalized on giving an accurate R judgment and giving an accurate R or K judgment to a studied item, respectively. See the Statistical Analyses section for formulae and additional details.

Regarding source memory, healthy older adults typically show large deficits in source memory relative to young adults (for reviews, see Old & Naveh-Benjamin, 2008; Spencer & Raz, 1995). Surprisingly, there were no significant effects involving Age (nor any significant effects of Encoding condition) on the pRRem measure of source memory,p’s ≥ 0.222. Similar results were obtained when source memory accuracy was computed with the binning scheme used for the ERP analysis (pR_ERP_), p’s ≥ 0.187. A likely explanation for this null age effect on source, enlarged upon in the Discussion, is that we minimized age differences in source accuracy by only probing source memory for trials attracting R responses.

### Task-Cue ERPs

In this section, we report the results relevant to age differences in preSMEs – differences in ERPs elicited by the task cue as a function of subsequent source memory. The number of trials contributing to these effects are reported in the left half of Table 5. We reiterate that encoding trials that went on to be forgotten (i.e., trials that subsequently received an N response) were not included in the analysis due to an insufficient number of trials in a high number of participants (see EEG/ERP Analysis). Thus, we focus on examining preSMEs associated with accurate and inaccurate source memory decisions on correctly recognized studied words (i.e., subsequent recollection effects). The analysis of preSMEs focused on average amplitude in the 500-1000 ms, 1000-1500 ms and 1500-2000 ms time windows. Figure 3 shows the grand average ERPs elicited by the task cues from a representative electrode (F2).

**Figure 3.**
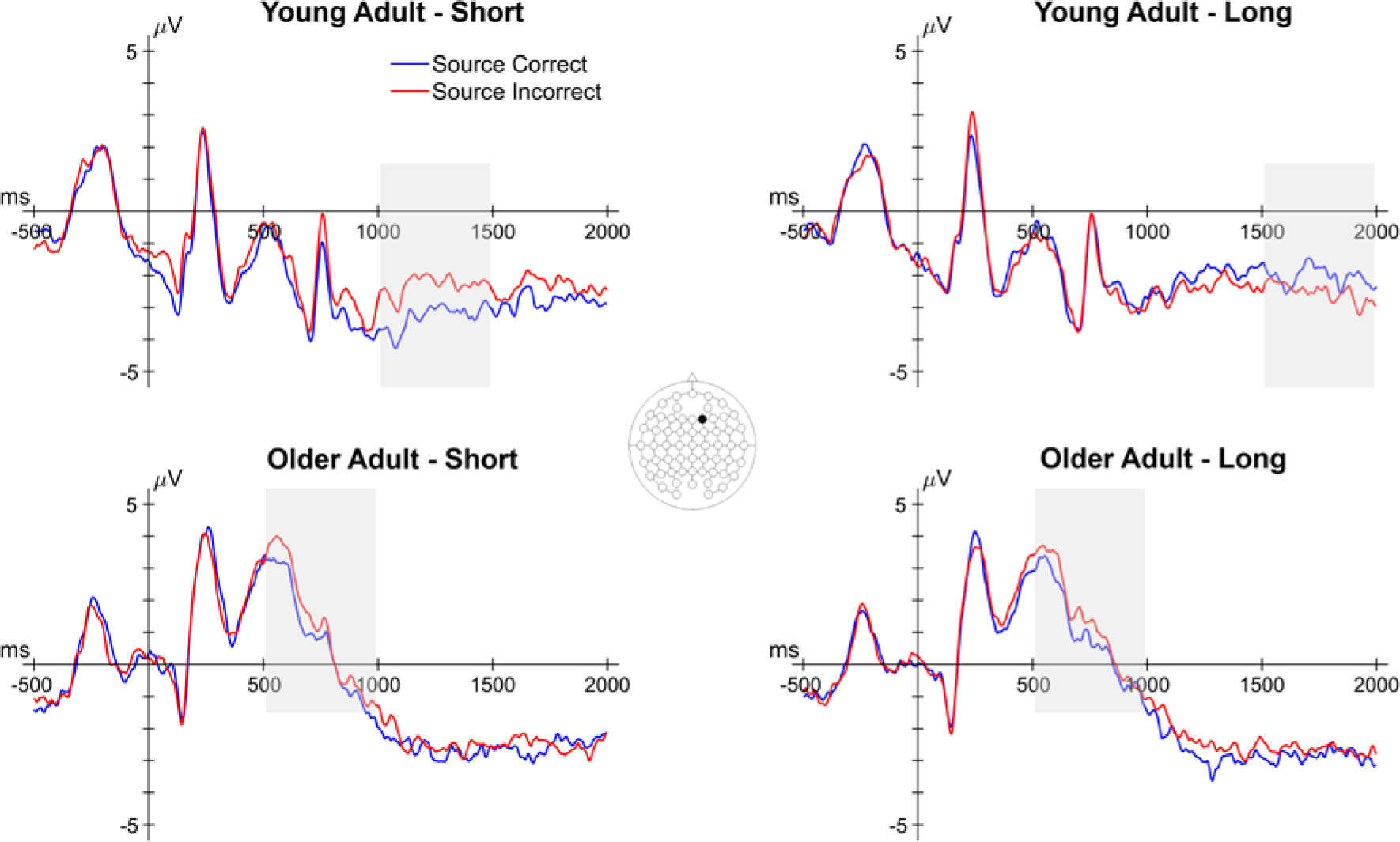
Grand average ERPs from a representative electrode (F2) of the task cue epoch for both encoding conditions in young and older adults. Shaded areas reflect time windows in which a significant effect involving subsequent memory was observed. Note that the early subsequent memory effect in older adults did not involve an interaction with encoding condition.

**Table 5.** Mean (and range) of the number of trials contributing to the bins for the task cue ERPs and word elicited ERPs as a function of age, encoding condition, and subsequent memory.

| Age Group | Encoding Condition | Task Cue ERPs |  | Study Word ERPs |  |
| --- | --- | --- | --- | --- | --- |
|  |  | Source Correct | Source Incorrect | Source Correct | Source Incorrect |
| Young | Short | 49<br>(21-81) | 45<br>(28-73) | 48<br>(21-83) | 46<br>(28-76) |
|  | Long | 47<br>(22-80) | 47<br>(31-76) | 49<br>(22-82) | 45<br>(29-73) |
| Old | Short | 42<br>(23-69) | 45<br>(20-80) | 42<br>(24-72) | 46<br>(16-76) |
|  | Long | 42<br>(24-72) | 46<br>(16-76) | 43<br>(23-68) | 44<br>(21-81) |

Note. The mean number of trials was rounded down to the nearest whole number.

#### 500-1000 ms

The omnibus ANOVA produced a significant four-way interaction between Memory, Age, AP Chain, and Laterality (see top panel of Table 6). A visual inspection of the 500-1000 ms time window (see Figure 4) indicates that older adults, but not young adults, show negative going (source correct < source incorrect) preSMEs in frontal electrodes. This interaction was decomposed with separate ANOVAs for each age group with the factors of Memory, AP Chain, Laterality, and Hemisphere, after collapsing across the two encoding conditions. No significant effects involving the Memory factor were observed in young adults, p’s involving Memory ≥ 0.124. In older adults, there was a significant interaction between Memory and AP Chain, F(2.19,50.37) = 4.54, *MSe* = 0.96, *p* = 0.013, partial-η^2^ = 0.17. Our expectation was that no preSMEs would be detectable in this early time window, thus finding this early effect in older adults was unexpected. Given this, we examined simple effects of subsequent source memory at each level of the AP chain factor (collapsed across other factors). A significant negative preSME was found for electrodes in the Fp chain, t(42.94) = 2.26, SEdiff = 0.18, *p* = 0.029, and F chain, t(42.94)= 2.04, SEdiff = 0.18, *p* = 0.048. No significant effects were observed at electrodes in the C, P, or O chains, t’s(42.94) < .80, SEdiff = 0.18, p’s ≥ 0.431.

**Table 6.** Results from the omnibus ANOVA for each time window in the task cue epoch.

| Model Term | *df* | *F* | *MS<sub>e</sub>* | *p* | $\eta_p^2$ |
| --- | --- | --- | --- | --- | --- |
| **500-1000 ms** |  |  |  |  |  |
| Memory X Age X AP X Laterality | 2.82,129.82 | 3.53 | 0.27 | 0.019 | 0.07 |
| **1000-1500 ms** |  |  |  |  |  |
| Memory X Age X Encoding X AP Chain | 1.87,85.93 | 4.43 | 2.99 | 0.017 | 0.09 |
| Memory X Encoding X Laterality | 1,46 | 4.79 | 0.48 | 0.034 | 0.09 |
| Memory X Encoding X AP Chain X Hemisphere X<br>Laterality | 3.32,152.59 | 2.70 | 0.10 | 0.042 | 0.06 |
| **1500-2000 ms** |  |  |  |  |  |
| Memory X Age X Encoding | 1,46 | 6.32 | 7.03 | 0.016 | 0.12 |
| Memory X Age X Encoding X Laterality | 1,46 | 4.82 | 0.54 | 0.033 | 0.09 |

*Note.* Only significant effects involving the Memory factor are reported. Reported degrees of freedom (*df*) are Greenhouse-Geisser corrected.

**Figure 4.**
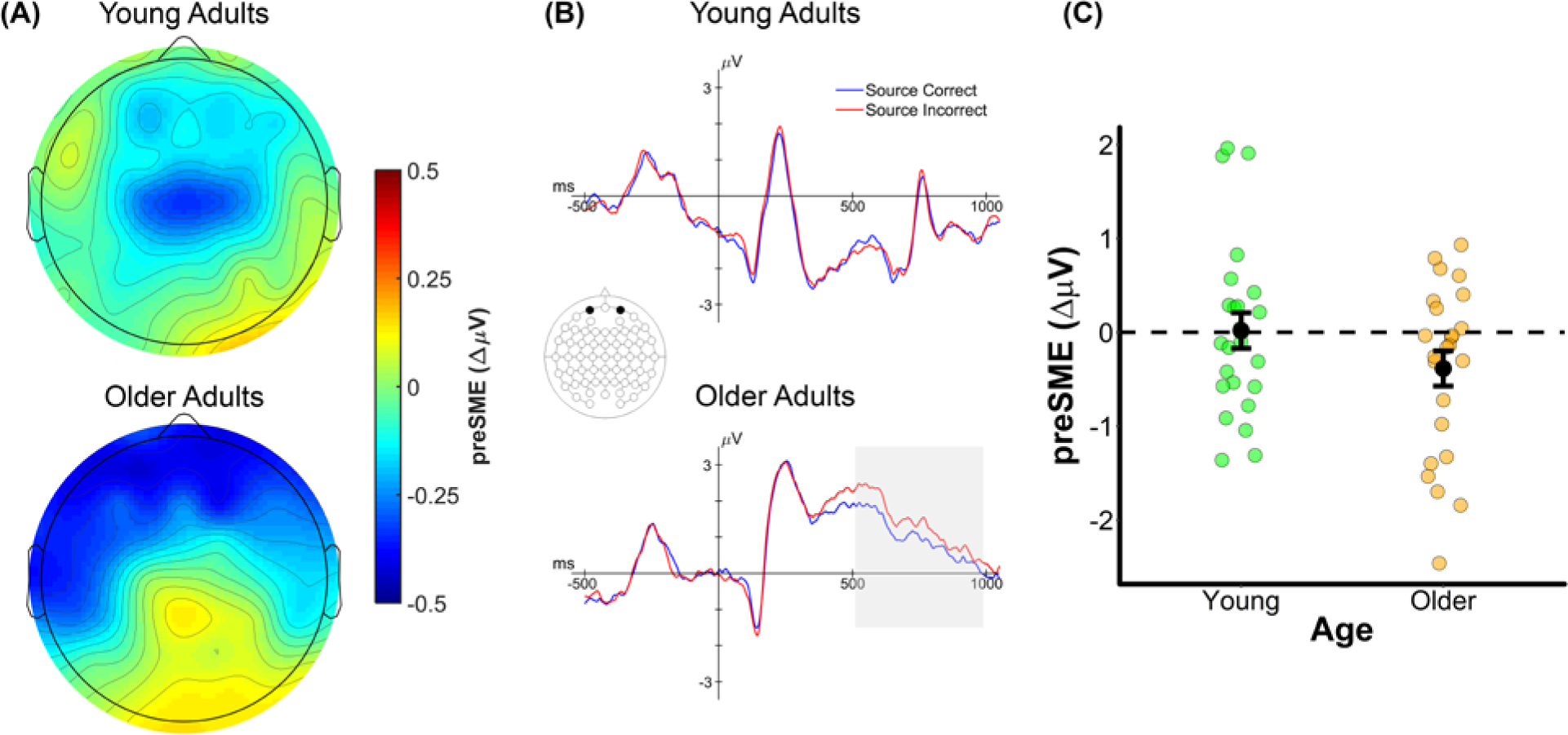
(A) Scalp plot of the preSMEs (source correct – source incorrect) in young and older adults in the 500-1000 ms time window following onset of the task cue (collapsed across encoding condition). Note the common scale used to depict the magnitude of the preSMEs in each age group. (B) ERP grand average traces from a virtual electrode created by averaging across electrodes Fp1 and Fp2. The ERP trace only plots the first 1000 ms the task cue epoch to highlight this early effect. (C) Plot showing the group means (black points) and individual preSMEs (green and orange points) for young and older adults. Data were average across electrodes Fp1 and Fp2 in the 500 to 1000 ms time window. Error bars reflect ±1 standard error of the mean.

#### 1000-1500 ms

The omnibus ANOVA for the 1000-1500 ms time window revealed three significant interactions involving a mixture of Age, Memory, Encoding Condition and the electrode site factors (see middle panel of Table 6). Together, these interactions indicate that the preSMEs in the 1000-1500 ms time window differed by both age group and encoding condition. Visual inspection of the ERP data (see Figures 3 and 5) suggests that the driver of these interactions is a negative going preSME at frontal electrodes for the short encoding condition in young but not older adults, with no corresponding preSME in the long encoding condition in either age group.

To test if age differences in preSMEs were present in both encoding conditions, the data from the short and long conditions were submitted to separate ANOVAs. The ANOVA for the short encoding condition revealed a significant interaction involving Memory, Age, AP Chain, Laterality, and Hemisphere, F(2.58,118.77) = 3.31, *MSe* = 0.38, *p* = 0.028, partial-η^2^ = 0.07. A further ANOVA on the data from the young group only revealed a significant interaction involving Memory, AP Chain, and Laterality, F(2.72,62.60) = 3.79, *MSe* = 0.43, *p* = 0.017, partial-η^2^ = 0.14; the analogous ANOVA for the older adults revealed no effects involving the Memory factor (all p’s ≥ 0.16). Post-hoc contrasts in young adults on the average ERP amplitudes from electrodes Fp1, Fp2, F1, and F2 showed a significant effect of subsequent memory, t(39.35) = 2.80, SEdiff = 0.29, *p* = 0.008 (see Figure 5C).

**Figure 5.**
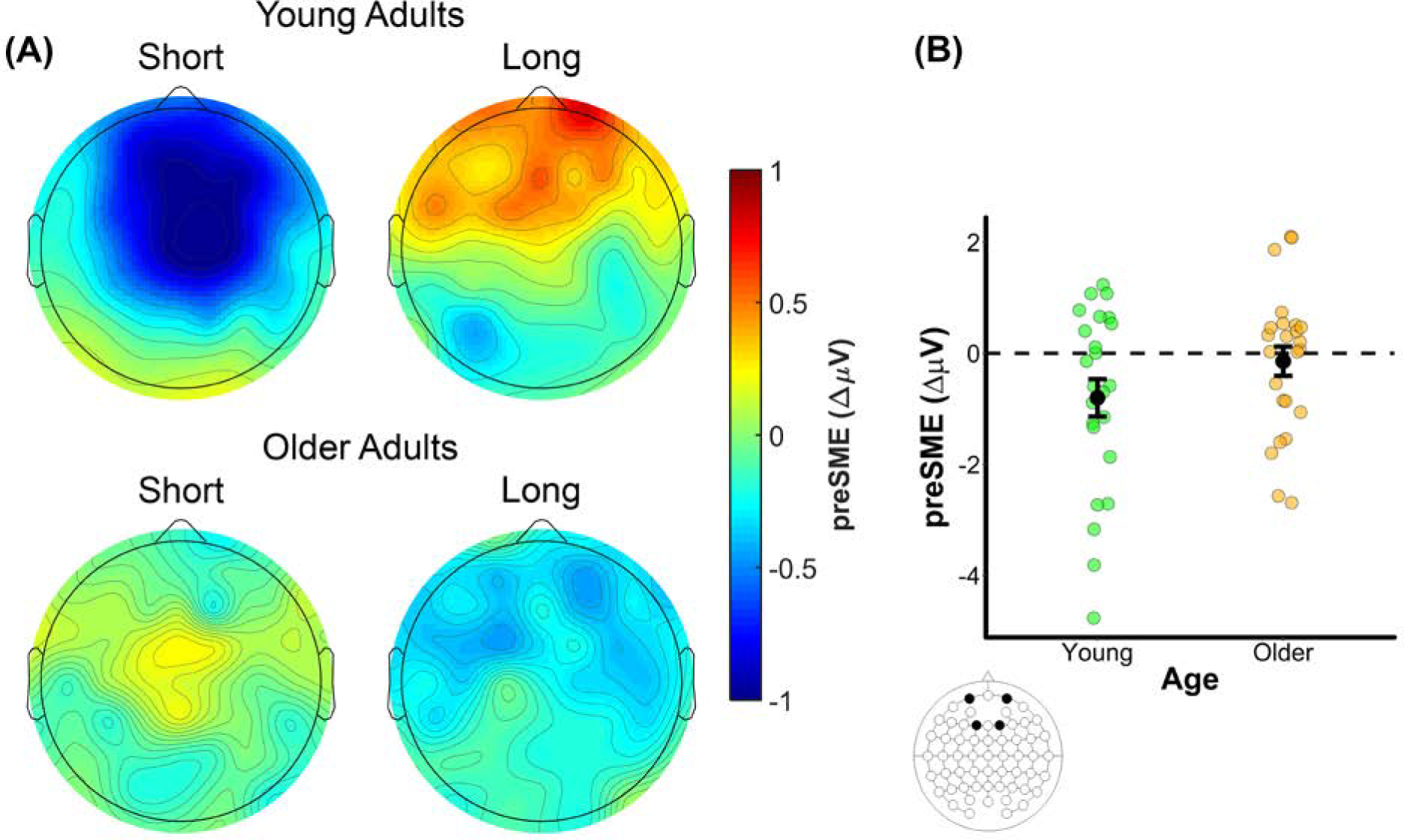
(A) Scalp plot of the preSMEs (source correct – source incorrect) in young and older adults in the 1000 to 1500 ms time window following onset of the task cue. Note the common scale used to depict the magnitude of the preSMEs in each age group and encoding condition. (B) Plot showing the group means (black points) and individual preSMEs (green and orange points) for young and older adults in the short encoding condition. Data were averaged across electrodes Fp1, Fp2, F1, and F2 in the 1000 to 1500 ms time window. Error bars reflect ±1 standard error of the mean.

A Memory, Age, AP Chain, Laterality, and Hemisphere ANOVA for the long encoding condition produced no effects of Memory or any interactions involving Age and Memory, all p’s involving Memory ≥ 0.15.

#### 1500-2000 ms

The omnibus ANOVA on the preSMEs in the 1500-2000 ms time window produced two significant interactions involving a combination of the factors of Age, Memory, and Encoding Condition (see bottom panel of Table 6). These interactions indicated that the preSMEs in the 1500-2000 ms time window differed both by age group and encoding condition. Contrary to what was reported above for the 1000-1500 ms time window, a visual inspection of the data suggests that the pattern of results in this later time window is driven by larger age differences in the long encoding condition relative to the short encoding condition (see Figures 3 and 6). Specifically, preSMEs appear to be positive-going in the long encoding condition for young adults but absent for older adults. Moreover, the preSMEs in the short condition for young adults, while still negative-going, appear to be smaller than those in the 1000-1500 ms time window.

The above interactions were decomposed with separate ANOVAs for the short and long encoding conditions. No significant effects involving Memory were observed in the ANOVA of the data from the short encoding condition (p’s ≥ 0.086). In contrast, there was a significant interaction involving Age and Memory in the long encoding condition, F(1,46) = 4.11, *MSe* = 9.86, *p* = 0.048, partial-η^2^ = 0.08. These results indicate that age differences in preSMEs are largest in the long encoding condition for the 1500-2000 ms time window (see Figure 6). Follow up group-wise ANOVAs revealed a significant effect of Memory in young adults, F(1,23) = 4.65, *MSe* = 8.10, *p* = 0.042, partial-η^2^ = 0.17, but not in older adults, all p’s involving Memory ≥ 0.234. Post-hoc contrasts conducted in young adults on the average ERP amplitudes from electrodes Fp1, Fp2, F1, and F2 revealed a significant effect of subsequent memory in the long encoding condition, t(56.96) = 2.46, SEdiff = 0.23, *p* = 0.017 (see Figure 6C).

**Figure 6.**
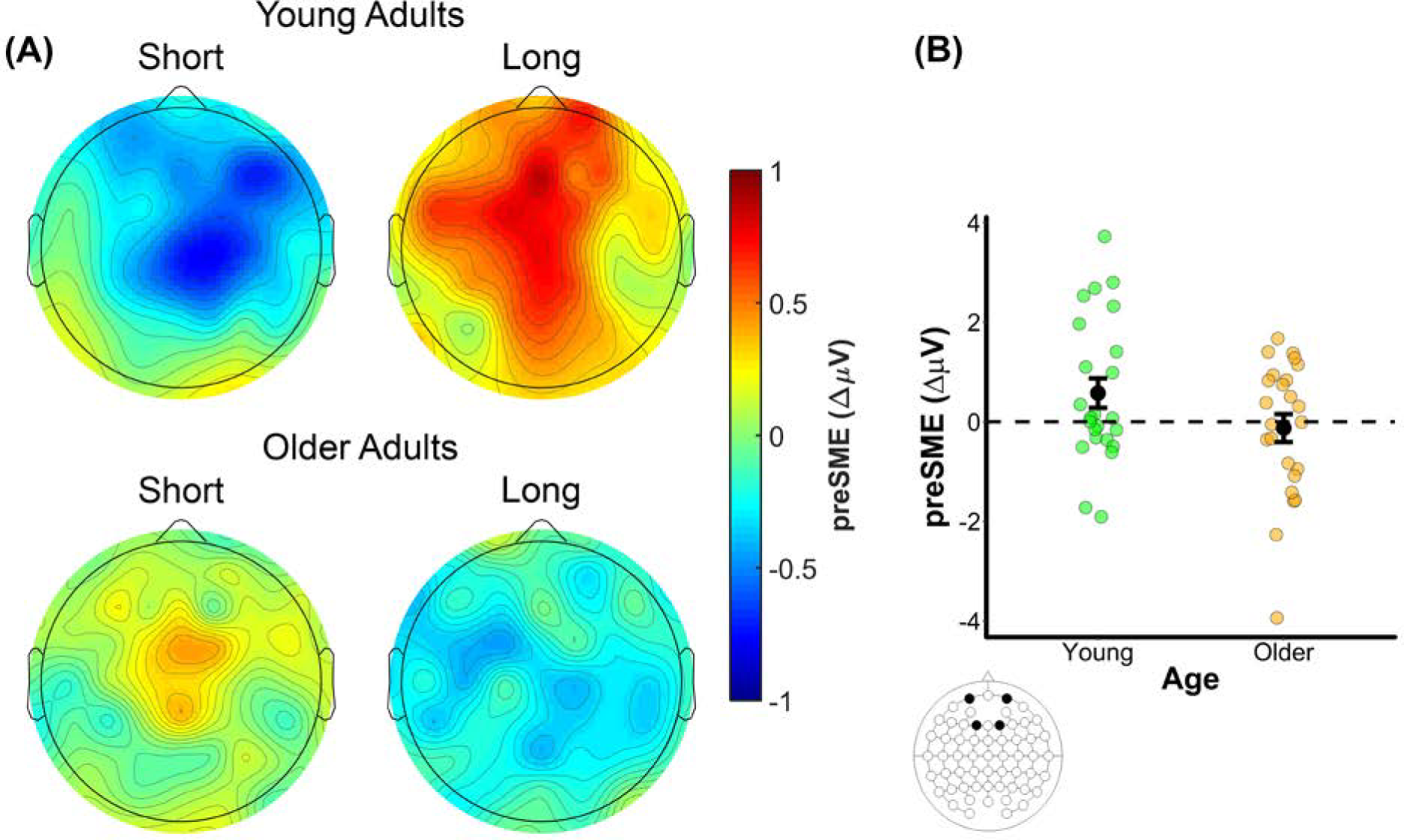
(A) Scalp plot of the preSMEs (source correct – source incorrect) in young and older adults in the 1500 to 2000 ms time window following onset of the task cue. Note the common scale used to depict the magnitude of the preSMEs in each age group and encoding condition. (B) Plot showing the group means (black points) and individual preSMEs (green and orange points) for young and older adults in the long encoding condition. Data were averaged across electrodes Fp1, Fp2, F1, and F2 in the 1500 to 2000 ms time window. Error bars reflect ±1 standard error of the mean.

#### Summary

Prestimulus SMEs elicited by the task cues demonstrated clear age differences. In the time windows typically associated with preSMEs in previous studies (the 1000-1500 ms and 1500-2000 ms time windows), young adults showed reliable preSMEs in both the short and long encoding conditions. Critically, there were qualitative differences in the preSMEs observed in young adults: whereas the preSME in the short encoding condition was negative (source correct < source incorrect) and was maximal in the 1000-1500 ms time window, the effect in the long encoding condition was positive and maximal in the 1500-2000 ms time window. Although we failed to detect preSMEs in older adults in either of the above time windows, preSMEs were apparent in the early (500-1000 ms) time window and invariant across the two encoding conditions.

### Study Word ERPs

In this section, we report the results relevant to age differences in SMEs – memory related differences in ERPs elicited by the study words. The numbers of trials contributing to this analysis are reported in the right half of Table 5, and grand average ERPs from a representative electrode (F2) are shown in Figure 7. The analyses focused on the average ERP amplitudes in the 300-600 ms, 600-1500 ms and 1500-2000 ms time windows and, as with the analysis of preSMEs, focused on subsequent recollection effects.

#### 300-600 ms

There were no significant effects involving the Memory factor in the omnibus ANOVA during the 300-600 ms time window, all p’s involving Memory ≥ 0.073.

**Figure 7.**
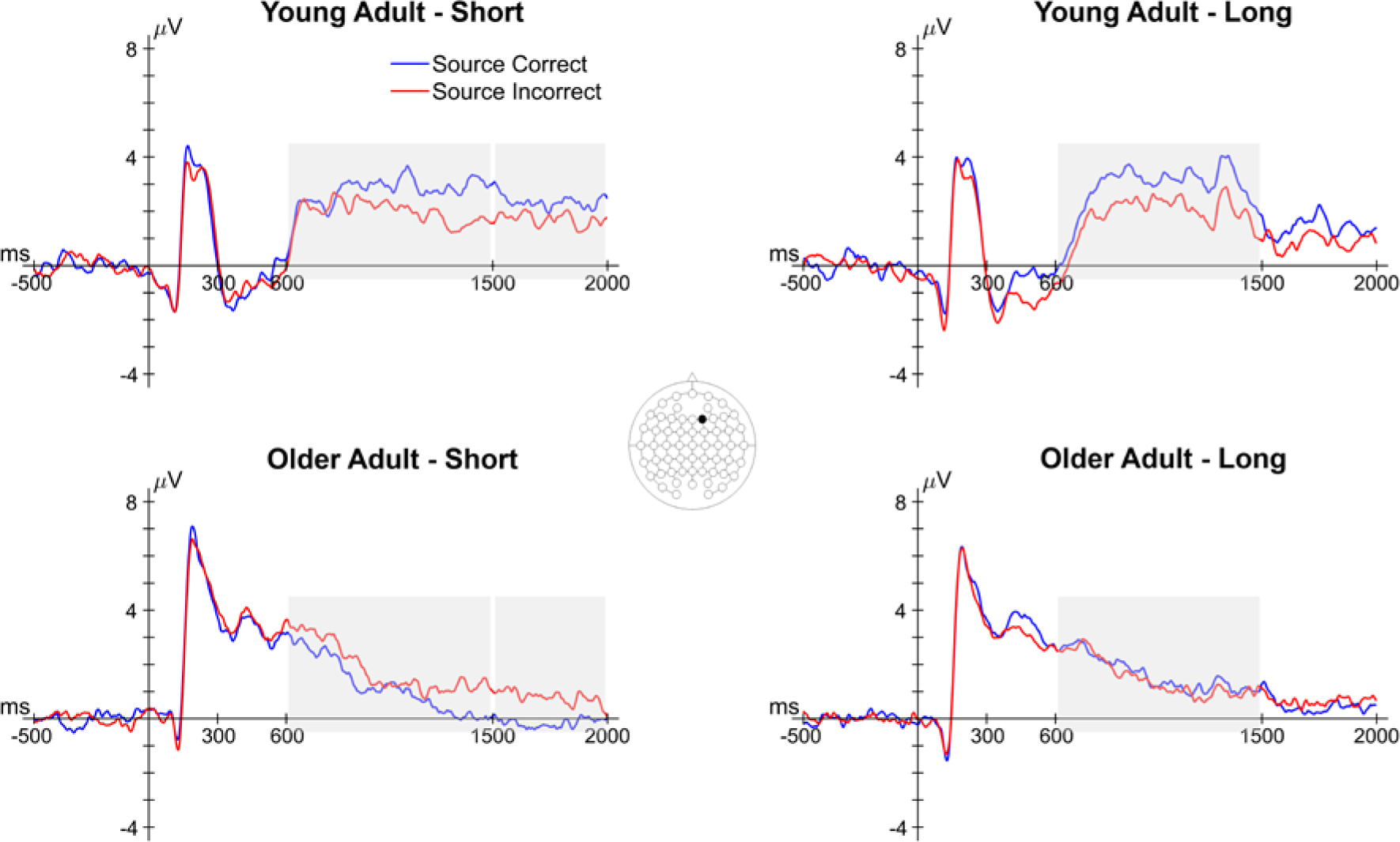
Grand average ERPs from a representative electrode (F2) of the study word epoch for both encoding conditions in young and older adults. Shaded areas reflect time windows in which a significant effect involving subsequent memory was observed. Note that the latter two time windows involved interactions between subsequent memory and age.

#### 600-1500 ms

The omnibus ANOVA of SMEs in the 600-1500 ms time window produced several interactions involving Memory, Age and the three electrode site factors (see middle panel of Table 7). Collectively, these interactions indicate that SMEs differed between young and older adults. A visual inspection of the data suggests that SMEs at mid-frontal electrodes were positive-going in younger adults and negative-going in older adults (see Figure 8A).

**Figure 8.**
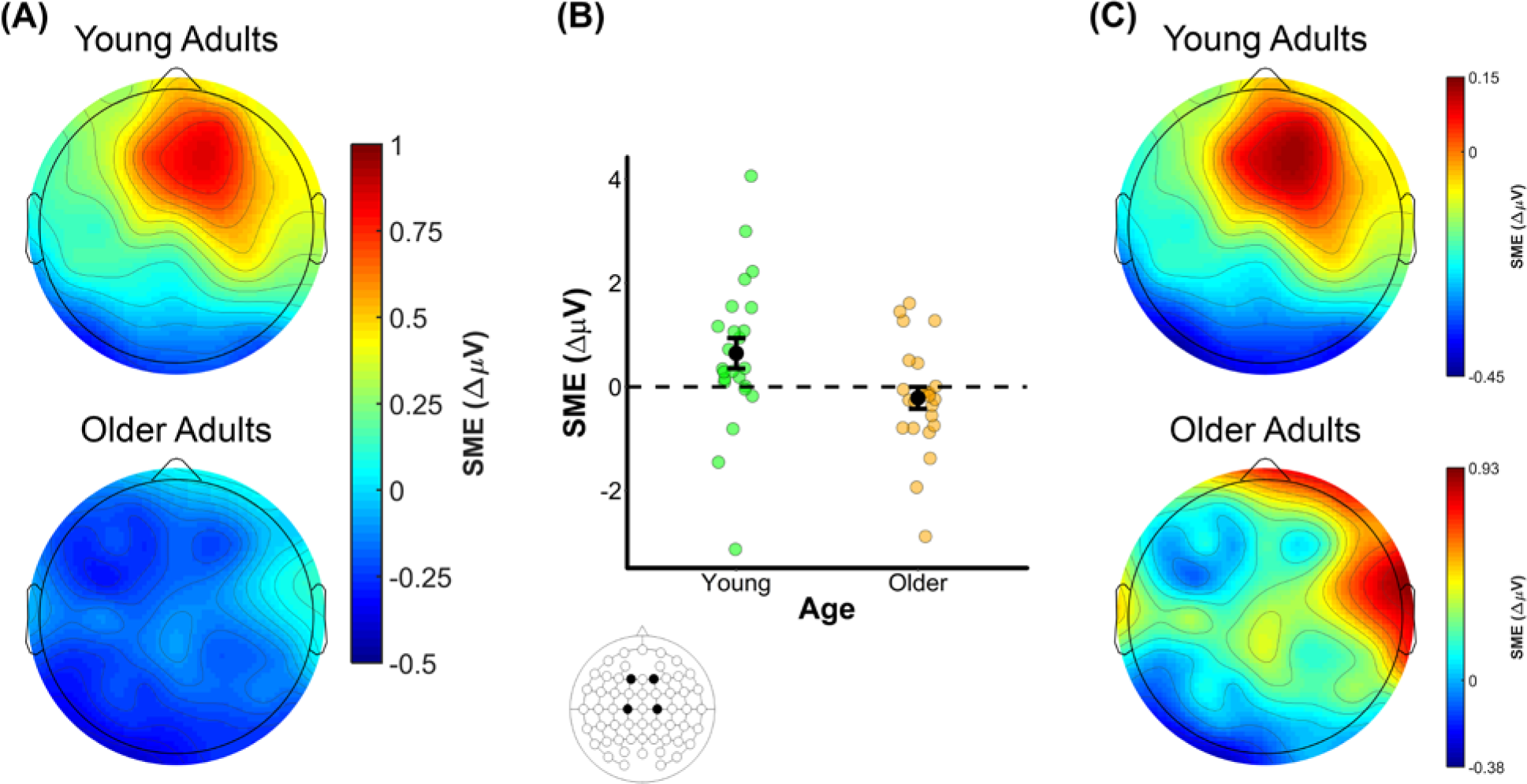
(A) Scalp plot of the SMEs (source correct – source incorrect) in young and older adults in the 600 to 1500 ms time window following onset of the study word. Note the common scale used to depict the magnitude of the preSMEs in each age group. (B) Plot showing the group means (black points) and individual SMEs (green and orange points) for young and older adults. Data were averaged across electrodes F1, F2, C1, and C2. Error bars reflect ±1 standard error of the mean. (C) Topography of the SMEs in the 600-1500 ms time window following onset of the study words (collapsed across encoding condition). Unlike the scalp map presented in (A), the scalp maps in this figure are min-max scaled separately for young and older adults to allow comparison of the topographies of the effects.

**Table 7.**
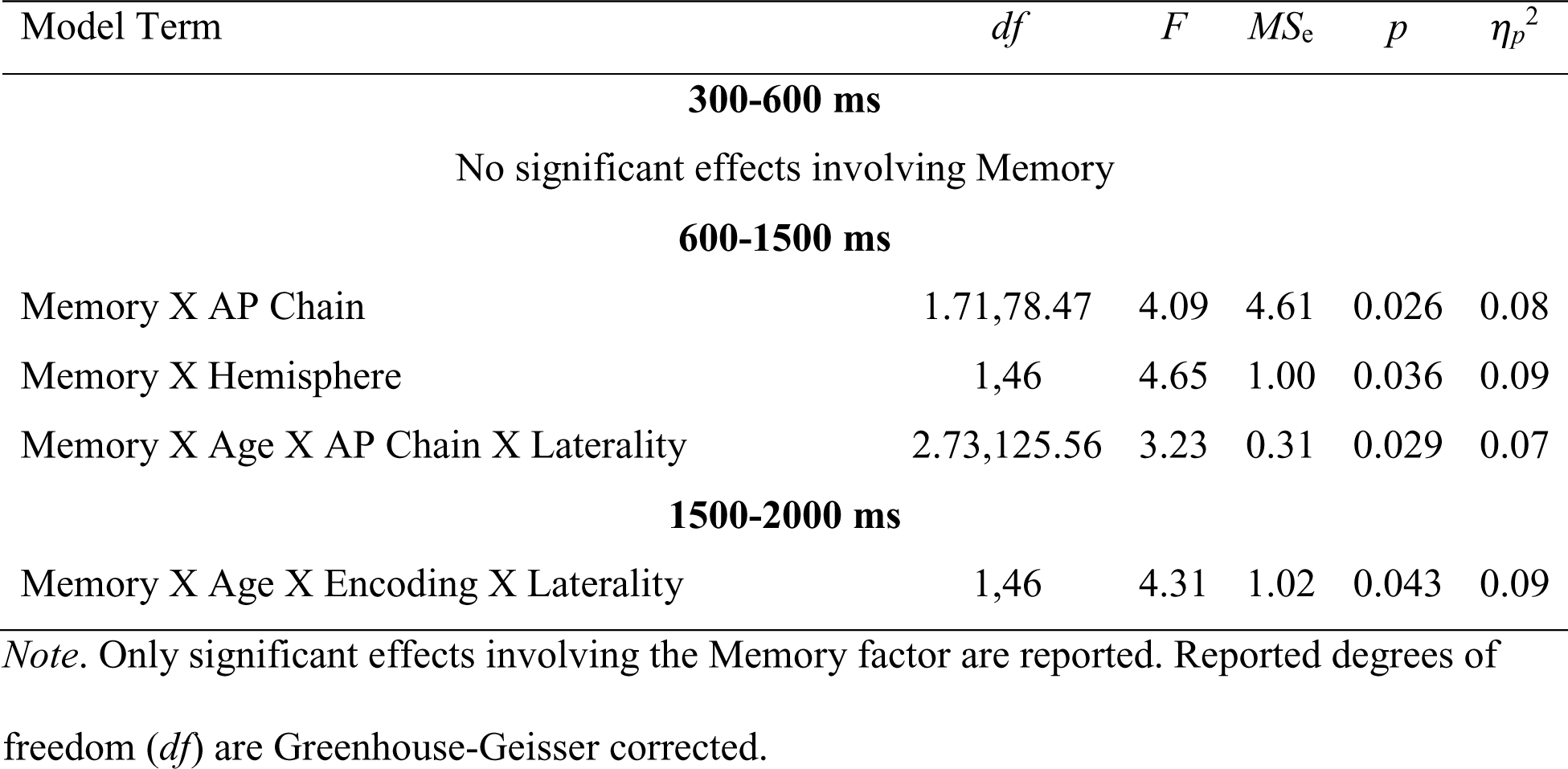
Results from the omnibus ANOVA for each time window in the study word epoch.

To examine whether SMEs were present in each age group, the young and older adult data (collapsed across encoding condition) were submitted to separate ANOVAs with the factors Memory, AP Chain, Laterality, and Hemisphere. These ANOVAs produced significant effects of Memory in both young and older adults. The ANOVA in young adults revealed an interaction involving Memory, AP Chain, and Laterality, F(2.86,65.68) = 3.64, *MSe* = 0.14, *p* = 0.019, partial-η^2^ = 0.14, and post hoc contrasts on the ERP amplitudes averaged over F1, F2, C1, and C2 revealed a significant effect of subsequent memory, t(37.22) = 3.14, SEdiff = 0.21, *p* = 0.003 (see Figure 8B). The ANOVA in older adults revealed a significant interaction between Memory and Hemisphere, F(1,23) = 4.46, *MSe* = 0.24, *p* = 0.046, partial-η^2^ = 0.16. However, the post hoc contrast examining a simple subsequent memory effect at electrodes F1, F2, C1, and C2 was not significant, t(39.33) = 1.26, SEdiff = 0.17, *p* = 0.215. Visual inspection of the data in older adults suggests that the interaction between Memory and Hemisphere was driven by relatively more negative SMEs in left hemisphere electrodes.

Given the significant interactions described above involving Memory and at least one electrode site factor in both young and older adults, it is possible that there are age differences in the configurations of the neural generators of the observed SMEs. To address this issue, we range normalized the SMEs using the procedure outlined by McCarthy and Wood (1985), and submitted these values to an ANOVA with the factors Age, AP Chain, Laterality, and Hemisphere (see Figure 8C). Note that the range normalization was conducted after collapsing across the two encoding conditions. This ANOVA did not produce any significant effects involving age, with all p’s involving Age ≥ 0.198, indicating that the scalp topographies of SMEs in young and older adults did not significantly differ.

#### 1500-2000 ms

The omnibus ANOVA for the SME in the 1500-2000 ms time window produced a significant four-way interaction between the Memory, Age, Encoding and Laterality factors (see bottom panel of Table 7). Visual inspection of the data (see Figure 9) suggests that the interaction was primarily driven by SMEs in young and older adults with opposing polarities in the short encoding condition.

**Figure 9.**
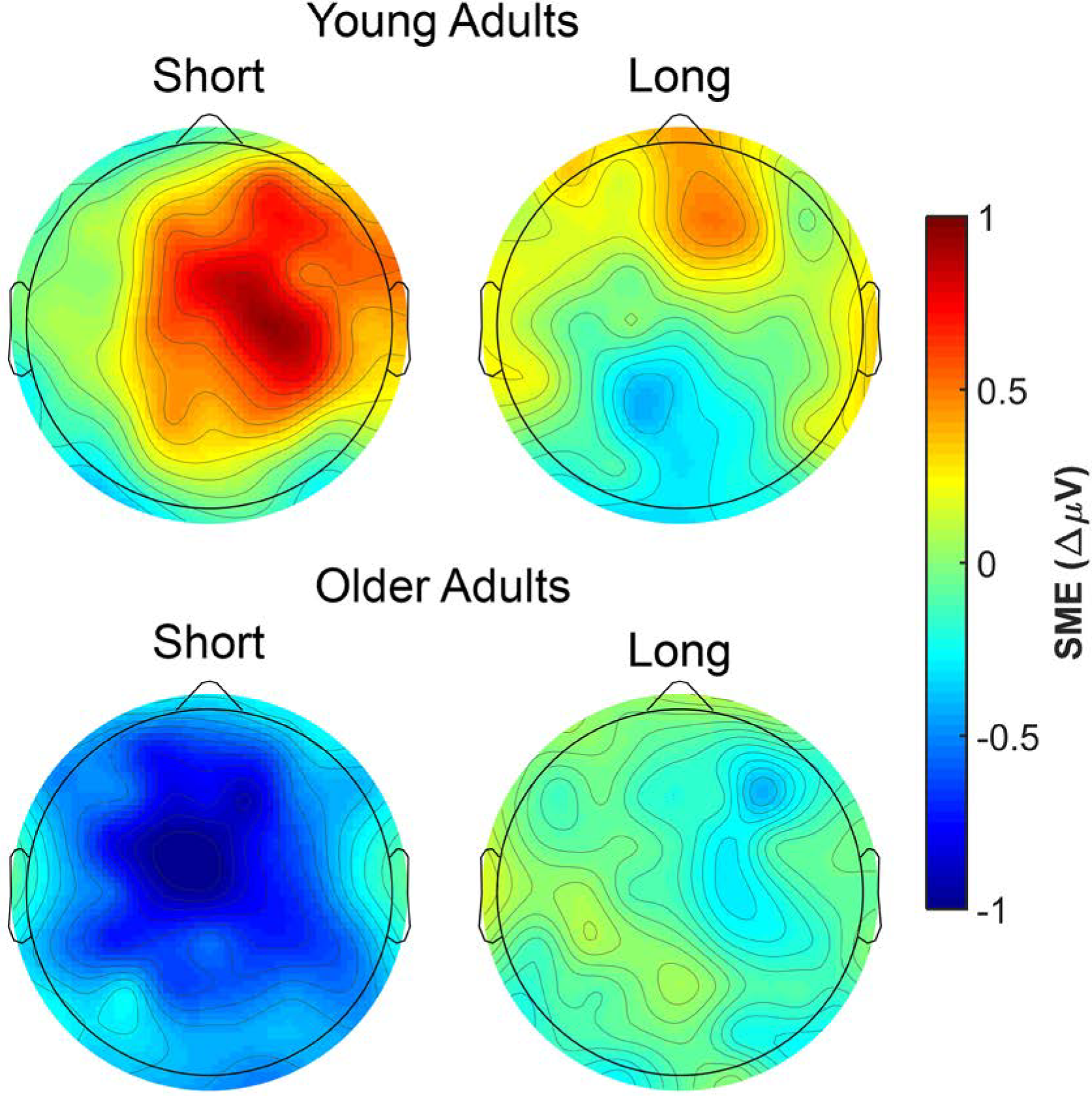
Scalp plot of the SMEs (source correct – source incorrect) in young and older adults for both encoding conditions in the 1500 to 2000 ms time window following onset of the study word. Note the common scale used to depict the magnitude of the SMEs in each age group.

A follow-up ANOVA with the factors Memory, Age, AP Chain, Laterality, and Hemisphere in the long encoding condition produced no effects involving Memory, all p’s involving Memory ≥ 0.213. In contrast, the analogous ANOVA conducted on data from the short encoding condition revealed an interaction between Age, AP Chain, Laterality, and Hemisphere, F(3.00,137.52) = 3.53, *MSe* = 0.45, *p* = 0.017, partial-η^2^ = 0.07. Follow-up ANOVAs produced a significant interaction between Memory and Laterality in young adults, F(1,23) = 4.66, *MS_e_* = 0.90, *p* = 0.042, partial-η^2^ = 0.07, but no significant effects involving the Memory factor in older adults, all p’s involving Memory ≥ 0.051. A post hoc contrast on the ERP amplitude averaged over electrodes F1, F2, C1, and C2 for the short encoding condition in young adults revealed a significant effect of subsequent memory, t(45.40) = 2.35, SE_diff_ = 0.29, *p* = 0.023.

#### Summary

Study words elicited robust SMEs in young adults, regardless of the encoding condition, that were maximal in the 600-1500 ms time window. Older adults showed reliable SMEs in the same time window, but their effects were reversed in polarity relative to young adults (i.e., source incorrect > source correct). A topographic analysis failed to detect age differences between young and older adults in the scalp topography of the SMEs in the 600-1500 ms time window. In the late time window, the SME in young adults persisted, but was only detectable for the short encoding condition.

## Discussion

The present study used ERPs to examine age differences in preparatory processes that are thought to benefit memory encoding (Otten et al., 2006; 2010). Specifically, preSMEs were examined in two encoding conditions assumed to differentially benefit from preparatory processing (see Introduction). There were several important findings. First, young, but not older, adults showed reliable preSMEs during the time window during preSMEs are typically observed (i.e., one second immediately prior to presentation of a study item). Second, most prior studies linking prestimulus neural activity to subsequent recollection have used either the Remember/Know task (e.g., de Chastelaine & Rugg, 2015; Otten et al., 2006; 2010; Gruber & Otten, 2010; Park & Rugg, 2010) or associative recognition (Addante et al., 2015). Thus, to our knowledge this is the first report of preSMEs with recollection operationalized using subsequent source memory. Third, the polarity of the preSMEs associated with subsequent recollection in young adults differed between two encoding conditions, with the negative preSMEs observed in the short encoding condition while positive preSMEs were observed in the long encoding condition. Finally, although preSMEs in older adults were undetectable in the standard preSME time window, we identified a preSME in an earlier window that was invariant across encoding conditions. Below, we discuss the implications of these and other findings.

### Behavioral Results

Although age differences in estimates of recollection and familiarity derived from R/K/N judgments were consistent with the prior literature (Koen & Yonelinas, 2014; Schoemaker et al., 2014), there were no such differences in estimates of source memory (i.e., memory for the encoding task). This null finding was unexpected given the extensive literature documenting sizeable age differences in source memory (Old & Naveh-Benjamin, 2008; Spencer & Raz, 1995), and particularly so in light of findings showing larger age-related differences in source memory than in R/K/N estimates of recollection (Duarte et al., 2008; for similar results, see Boywitt, Kuhlmann, & Meiser, 2012; Kuhlmann & Boywitt, 2016). While there are several possible explanations for the present findings, a prime candidate is that the null effects for source memory arose because the judgments were limited to trials that had been endorsed with an R response. That is, source memory was only assessed for those trials for which participants reported a recollective experience. This might have resulted in participants constraining R responses only to those trials for which they were confident they could recollect the source detail (cf., Parks, 2007). By this account, the coupling of significant age differences in R/K/N estimates of recollection with null effects of age on source memory accuracy reflects the fact that older adults recalled source details less frequently than young adults.

### Prestimulus Subsequent Memory Effects in Young Adults

An important finding in the young adults’ ERPs concerns the different pattern of preSMEs in the two encoding conditions. The preSME in paradigms similar to ours (e.g., Otten et al., 2006) typically takes the form of a negative going effect for subsequently recollected versus non-recollected study events that is maximal at frontal midline electrodes. Prior preSMEs are usually largest during the second or so immediately preceding the onset of the study item (Galli et al., 2012; Galli et al., 2013; Otten et al., 2006; 2010). In the present young adult sample, preSMEs in the short encoding condition exhibited this characteristic pattern: a lower ERP amplitude for source correct trials relative to source incorrect trials that was maximal in the 1000-1500 ms time window following onset of the task cue. A strikingly different pattern was observed in the long encoding condition, however; here, preSMEs demonstrated a *positive* effect that was maximal in the 1500-2000 ms time window. The present findings add to prior reports of polarity reversals in preSMEs (de Chastelaine & Rugg, 2015; Padovani et al., 2011; for similar findings, see Galli, Wolpe, & Otten, 2011; Gruber & Otten, 2010). Critically, the polarity reversals observed in these prior studies were when the task cues signaled different task requirements: semantic versus phonological tasks in de Chastelaine and Rugg (2015), and semantic versus emotional in Padovani et al. (2011). Here, however, the polarity reversal in preSMEs were a consequence of an encoding manipulation (i.e., the presentation duration of the study items) that held the nature of study task constant.

We do not have a ready explanation for the polarity reversal observed in the present study. To the extent that the cross-over interaction (see Figures 5 and 6) is replicable, these findings presumably indicate that qualitatively different neural processes, and likely qualitatively different preparatory processes, were engaged in each encoding condition. It is plausible that the interaction reflects the greater incentive for engaging preparatory processing that accompanied the short encoding condition (see Introduction). Notably, the stimulus durations employed in prior ERP studies of preSMEs that employed visual stimuli (Galli et al., 2012; Galli et al., 2013; Gruber & Otten, 2010; Otten et al., 2006; 2010; Padovani et al., 2011; Padovani et al., 2013) in no case exceeded 500 ms. Together with the present findings, this observation suggests that negative-going preSMEs occur when the study conditions place a particularly high premium on prestimulus preparation. Yet, this explanation obviously does not provide an account for the finding that preSMEs in the long encoding condition were reliably *reversed* in polarity, rather than merely attenuated or non-existent. Clearly, additional research is required to identify the boundary conditions for preSMEs in ERPs.

Despite its opaque functional significance, the polarity reversal in preSMEs observed here between the short and long encoding conditions bears on current theoretical accounts of these effects. As discussed in the Introduction, one account is that the effects reflect spontaneous neural fluctuations or states that are differentially conducive to successful encoding (e.g., Ezzyat et al., 2017; Guderian et al., 2009; Fernandez et al., 1999; Yoo et al., 2012; Fell et al., 2011). The present results, however, are apparently inconsistent with this account, since it would seem to predict a null effect for the encoding duration manipulation on the magnitude or direction of preSMEs. Other things being equal, spontaneous neural fluctuations or states that modulate likelihood of successful encoding should exert an influence regardless of other factors. Of course, it is possible that other things are not equal, and that spontaneous neural flucations modulate memory encoding in a context-dependent manner. Current evidence does not allow the likelihood of this possibility to be assessed.

A second account proposes that preSMEs reflect the engagement of task-specific preparatory processes (e.g., adopting a ‘semantic’ task set) that enhance subsequent encoding (de Chastelaine & Rugg, 2015; Otten et al., 2006; 2010). An important tenet of this account is that the task cue carries information that allows the appropriate task set to be proactively engaged. This account implies that preSMEs might vary according to the nature of the study task (e.g., whether it is semantic or phonological, cf. de Chastelaine & Rugg, 2015; Otten et al., 2006), but has nothing to say about other factors that might influence the effects. The present findings indicate that such additional factors must exist: whereas the short and long study durations led to differences in task difficulty (indicated by reliable RT differences), as already noted, the two conditions utilized identical task cues and study judgments. Thus, our observation of condition-dependent differences in the polarities of the associated preSMEs indicates that these effects reflect more than the adoption of a specific (in this case, semantic) task set.

### Age Differences in Prestimulus Subsequent Memory Effects

In older adults, no preSMEs were detectable in the second or so just prior to the onset of the study item. This finding is in line with our hypothesis that preSMEs in older adults would be reduced in comparison to those in young adults, and is consistent with prior research indicating that older adults have a reduced capacity to deploy proactive processes in anticipation of an upcoming task (Braver et al., 2001; Braver et al., 2009; Duverne et al., 2009; Morcom & Rugg, 2004). However, this finding is insufficient to conclude that older adults are *incapable* of engaging preparatory processes thought to benefit memory encoding. This conclusion will need to await evidence that older adults consistently fail to demonstrate preSMEs across a wide array of tasks, as well as when neural activity is assessed with measures other than ERPs. Notably, studies using fMRI and intracranial recordings have identified robust preSMEs in brain regions – such as the hippocampus - whose neural activity is essentially inaccessible to scalp EEG recordings (e.g., Adcock et al., 2006; de Chastelaine & Rugg, 2015; Fell et al., 2011; Park & Rugg, 2010).

We did however observe preSMEs in older adults during an earlier time window (500-1000 ms following onset of the task cue). These effects, which were insensitive to the study duration manipulation, took the form of an enhanced positive-going deflection elicited by task cues preceding study items associated with inaccurate source memory judgments (see Figure 4). Intriguingly, the scalp distribution of this effect is somewhat reminiscent of the well-studied ‘P3a‘ component, which is widely considered to be a correlate of attentional capture or novelty detection (for review, see Soltani & Knight, 2000; Polich, 2007, 2012). Therefore, it is tempting to speculate that the greater positivity of the ERPs elicited by task cues preceding study items for which recollection failed rather than succeeded reflects the potency of these cues in capturing attention, and a corresponding reduction in resources available for the encoding of the study item. We caution however that not only is this account highly speculative, but that the reliability and generality of the early preSMEs in older adults reported here remains to be established. Post-stimulus Subsequent Memory Effects

The robust SMEs that we observed in our young adults represent a replication of numerous prior reports (Friedman & Johnson, 2000; Rugg, 1995; Wilding & Ranganath, 2012). By contrast, and as predicted, SMEs in older participants were markedly attenuated; indeed, if anything, the effects were reversed in their polarity. These age differences are consistent with prior reports (e.g., Cansino et al., 2010; Friedman & Trott, 2000; Gutchess et al., 2007), and are in keeping with the proposal that aging is associated with a reduction in the efficacy of encoding processes that support successful recollection (for reviews, see Craik & Rose, 2012; Friedman & Johnson, 2014; Werkle-Bergner et al., 2006).

What might account for the age differences in SMEs observed here and in prior reports? The extensive literature on the effects of age on SMEs manifest in fMRI BOLD activity offers some intriguing clues. These effects take one of two forms (Kim, 2011; Rugg et al., 2015): ‘positive’ SMEs, where BOLD activity is greater for later remembered than forgotten items, and ‘negative’ SMEs, where BOLD activity is greater for later forgotten items. It has been reported in numerous studies that semantically elaborative study tasks are associated with positive SMEs predictive of later recollection in several cortical regions (most notably in lateral prefrontal cortex), as well as the medial temporal lobe, including the hippocampus (Eichenbaum, Yonelinas, & Ranganath, 2007; Kim, 2011; Yonelinas, Aly, Wang, & Koen, 2010). Negative SMEs, by contrast, are typically found in regions belonging to the ‘default mode network’. These regions include the posterior cingulate, lateral parietal cortex, and medial prefrontal cortex (e.g., Otten & Rugg, 2001; for review, see Kim, 2011). The balance of the evidence suggests that positive SMEs differ little, if at all, as a function of age (indeed, some effects are larger in older adults; e.g., de Chastelaine et al., 2016; de Chastelaine, Wang, Minton, Muftuler, & Rugg, 2011; for review, see Maillet & Rajah, 2014a). However, negative SMEs in default mode regions are consistently reported to be attenuated, or even reversed, in older adults (e.g., de Chastelaine, Mattson, Wang, Donley, & Rugg, 2015; Mattson et al., 2014; see Maillet & Rajah, 2014a). Viewing the present and previous (e.g., Friedman & Johnson, 2014) findings of age-related reductions in ERP SMEs through the lens of this fMRI evidence, it is possible that these effects reflect the same neural processes that are responsible for negative SMEs in fMRI data. Given the frontal distribution of the ERP SMEs that are typically observed in young adults, we conjecture that age-related reductions in these effects reflect differential encoding-related activity in the midline frontal regions manifesting negative SMEs in fMRI studies (e.g., Dennis et al., 2008; Leshikar & Duarte, 2014; Maillet & Rajah, 2014a; Morcom, Good, Frackowiak, & Rugg, 2003). Future research directly comparing ERP and fMRI data in the same samples of participants will be required to examine this possibility.

### Conclusions

The present results are consistent with the proposal that age-related reductions in memory performance are associated with reduced efficacy of the neural processes supporting memory encoding (Craik, 1983; Craik & Rose, 2012; Friedman & Johnson, 2014; Werkle-Bergner et al., 2006). Importantly, while the prior literature has mainly focused on age differences in encoding-related neural activity occurring shortly *after* the presentation of a study item, the present results indicate that age differences are also evident in the encoding-related neural activity that occurs in the second or so *before* a study item is experienced. Thus, explanations of age-related decline in the efficacy of episodic encoding must account for neurocognitive processes engaged both before and after a study event is encountered.

